# Curcumin-Magnesium complex loaded DNA hydrogels: concentration dependent swelling kinetics and selective cytotoxicity via Oxidative Stress induced apoptosis

**DOI:** 10.64898/2026.05.10.724072

**Authors:** Jugal Patil, Satyam Bhalerao, Ankur Singh, Geethu Prakash, Hatif Alam, Prachi Thareja, Dhiraj Bhatia

## Abstract

Curcumin is a naturally occurring polyphenol that demonstrates considerable anti-cancer activity, however the aqueous insolubility, rapid metabolism and relatively low bioavailability are limiting to its clinical application. As such, a curcumin-magnesium (Cur-Mg) coordination complex was synthesized and subsequently encapsulated within DNA hydrogels (Cur-Mg-Hgel). The Cur-Mg complex was fully characterized using UV-Vis spectroscopy, FTIR and X-ray diffraction (XRD). UV-Vis, FTIR and XRD all support the formation of a coordination complex and suggest a decreased level of crystallinity compared to free curcumin. DNA hydrogels were formed and characterized using atomic force microscopy, rheology and swelling kinetic studies. In vitro cytotoxicity studies utilizing an MTT assay demonstrate dose dependent inhibition of HeLa cell proliferation and a slightly better retention of RPE-1 viability at low concentrations (suggesting some difference in sensitivity) though significant cell death is seen at higher concentrations and both cells. Intracellular production of ROS was measured using the DCFH-DA assay and is seen to increase when HeLa cells are treated with Cur-Mg-Hgel in comparison to un-treated controls. Annexin V/PI staining demonstrates primarily late or early apoptotic activity with minimal necrosis following treatment with Cur-Mg-Hgel. The evidence presented strongly supports the notion that Cur-Mg-Hgel is a ROS-modulating, pro-apoptotic Hydrogel suitable for cancer treatment.

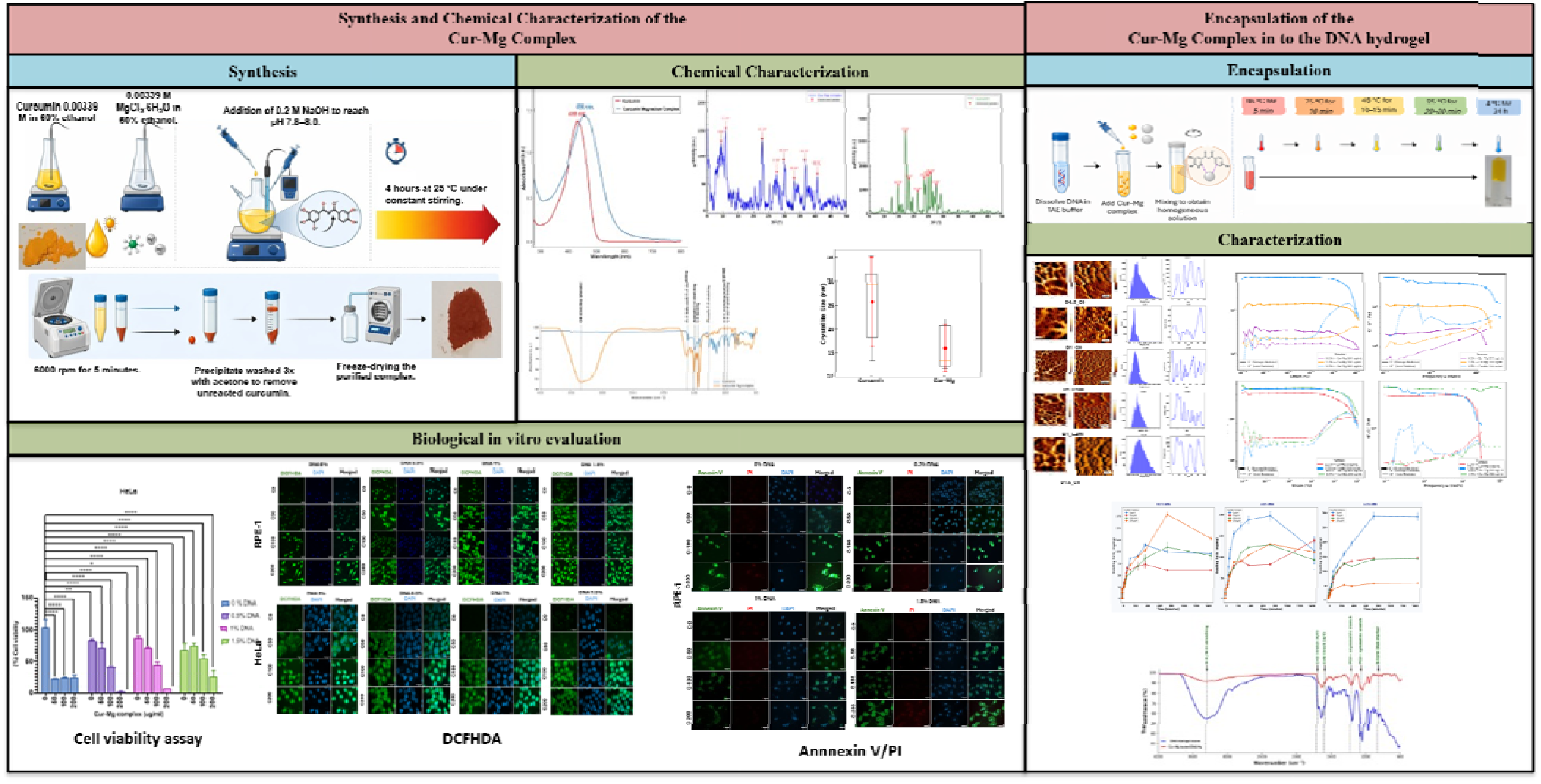

## 1. Introduction

Deoxyribonucleic acid (DNA) has emerged as a structurally programmable and biocompa ible scaffold for the engineering of next-generation drug delivery^1–9^, wound healing^7,10,11^, cell proliferation^4,12^ and biosensing^13,14^ platform owing to its predictable Watson-Crick base-pairing chemistry^9^, inherent sequence specificity and remarkable functional versatility^15^.Among the diverse architectures accessible through DNA nanotechnology^16,17^, DNA hydrogels three-dimensional^18^, hydrophilic polymeric networks composed of crosslinked DNA chains have attracted considerable interest as tunable therapeutic carriers capable of encapsulating a wide range of bioactive molecules, including small-molecule chemotherapeutics^19^, functional nucleic acids^20^, therapeutic proteins, and nanoparticles^15,21,22^. These structures can be fabricated through either chemical crosslinking strategies as some DNA nano structures are prepared in some cited studies^20,23–26^, employing enzyme-mediated ligation, photo crosslinking, or covalent DNA-polymer conjugation to yield mechanically robust and irreversible networks, or through physical crosslinking approaches exploiting hydrogen bonding, metal-ion coordination, hydrophobic interactions, and electrostatic forces to produce reversible, injectable, and minimally cytotoxic scaffolds^27,28^.In the context of cancer nanomedicine^4,29^, DNA hydrogels additionally offer susceptibility to nuclease-mediated degradation within the tumour microenvironment^30^, providing an inherent mechanism for localised, stimuli-responsive drug release, while surface functionalisation^31^ with tumour-targeting aptamers enables active receptor-mediated cellular internalisation collectively addressing the dual imperatives of therapeutic efficacy and systemic safety that continue to challenge conventional chemotherapy^32,33^.

Curcumin (1,7-bis(4-hydroxy-3-methoxyphenyl)-1,6-heptadiene-3,5-dione), is a polyphenolic natural product derived from rhizome of Curcuma longa, represents a paradigmatic example of a therapeutically promising yet clinically underperforming compound^34^. Its well-documented anticancer bioactivities encompass modulation of multiple oncogenic signalling pathways including NF-κB, PI3K/Akt/mTOR, Wnt/β-catenin, and MAPK cascades alongside potent antioxidant and anti-inflammatory properties^35^. Despite this compelling pharmacological profile Curcumin’s clinical translation has been severely hindered by its extreme aqueous insolubility, rapid metabolic degradation via two pathways which included glucuronidation and sulfation, and consequent poor oral bioavailability, which results in subtherapeutic systemic concentrations after conventional administration, despite this compelling pharmacological profile^36^. Through coordination at the β-diketone and phenolic hydroxyl moieties, metal complexation with divalent cations like magnesium (II) has been suggested as a logical method to improve the physicochemical profile of curcumin. This results in stable six-membered chelate rings that delocalize π-electrons, extend conjugation, and promote partial amorphization, a structural transition commonly linked to improved aqueous dissolution and bioavailability^37,38^. According to a study, magnesium is biologically ubiquitous divalent cation which act as pro-apoptotic ion for cervical cancer^39^. Integration of the curcumin–magnesium (Cur-Mg) coordination complex within a programmable DNA hydrogel matrix therefore represents a dual-engineering strategy that simultaneously addresses the biopharmaceutical limitations of curcumin.

The mechanistic basis underlying the preferential cytotoxicity of curcumin-based formulations toward malignant cells is intimately linked to the fundamentally distinct redox biology of cancer cells with respect to their normal counterparts^40^. Neoplastic transformation is characterised by a chronic state of elevated intracellular reactive oxygen species (ROS) production, arising from oncogene-driven metabolic reprogramming, mitochondrial dysfunction, increased oxidative phosphorylation, and heightened NADPH oxidase activity a biochemical hallmark that simultaneously drives tumorigenic signalling and renders cancer cells uniquely vulnerable to further pro-oxidant insults^35,41^. Specifically, cancer cells operate at a redox setpoint that is constitutively elevated relative to normal cells, yet this elevated basal ROS level approaches the threshold beyond which further oxidative stress becomes cytotoxic consequently, a relatively modest additional ROS burden readily achievable through pharmacological pro-oxidant intervention is sufficient to precipitate irreversible oxidative damage^41,42^. This concept of exploiting the differential redox threshold between malignant and non-malignant cells often termed oxidative stress-based selective anticancer therapy has been extensively validated across multiple cancer types, including cervical carcinoma, wherein HeLa cells exhibit markedly elevated baseline ROS relative to non-transformed epithelial cell lines^43^. Curcumin potentiates this vulnerability through a pleiotropic pro-oxidant mechanism that encompasses direct generation of superoxide and hydrogen peroxide via redox cycling of its phenolic moieties, inhibition of glutathione synthesis and thioredoxin reductase activity, downregulation of the Nrf2-mediated antioxidant response in cancer cells, and induction of mitochondrial ROS through disruption of Complex I/III of the electron transport chain mechanisms collectively amplified upon metal complexation, which modifies the electronic structure of the curcumin chromophore and may enhance its capacity to generate ROS via ligand-to-metal charge-transfer processes^44^.

In this study, we formulated a curcumin-magnesium(II) coordination complex encapsulated within DNA hydrogels of varying concentrations (0.5-1.5% w/v), forming a well-defined Cur-Mg-DNA hydrogel (Cur-Mg-Hgel) platform for delivery. DNA matrix provides the possibility of controlling the crosslinking density in a tunable manner with the change in Cur_Mg concentration due to the Mg mediated interactions and the architecture of the network formed in the resulting hydrogel, physical and chemical characteristic of the formed hydrogel analyzed by UV-Vis, FTIR, PXRD, AFM, rheology and swelling kinetics confirm that a progressive condensation of the gel takes place and gradual transition from Fickian to anomalous diffusion mechanism of Cur-Mg complex across the network of Cur-Mg-Hgel is achieved. Moreover, using MTT, ROS and apoptosis assays on HeLa and RPE-1 cells, an increased anticancer activity and enhanced specificity is exhibited. The work, therefore, combines metal coordination chemistry and DNA hydrogel engineering for redox mediated cancer therapy using a novel functional drug delivery platform

## 2. Results and Discussion

### 2.1 Synthesis and Chemical Characterization of the Cur-Mg Complex

The curcumin-magnesium (Cur-Mg) complex was prepared through a careful coordination technique Vizuallize in **Figure 1**. A equimolar of curcumin and MgCl_2_.6H_2_O was used. The curcumin was dispersed in 60% ethanol (0.00339 M) with slight heating (60°C) to get a clear yellow solution and the solution was cooled down to 25°C, keeping away from light. 0.00339 M of aqueous solution of MgCl_2_.6H_2_O was stirred at 400 rpm (25 1 C) and the curcumin solution was added drop by drop slowly. The solution’s pH was maintained to be around 7.8-8.0 using 0.2M NaOH, in order to achieve effective deprotonation of the - diketone moiety to facilitate chelation. During addition the yellow color changed to reddish-yellow which proves the formation of complex. This reaction was carried out for 4h at 25 C. The complex obtained was separated using centrifugation (6000 rpm, ∼4000 g, 5 min), washed with acetone to extract out unreacted curcumin, and then lyophilized to get a solid.

**Figure. 1:**
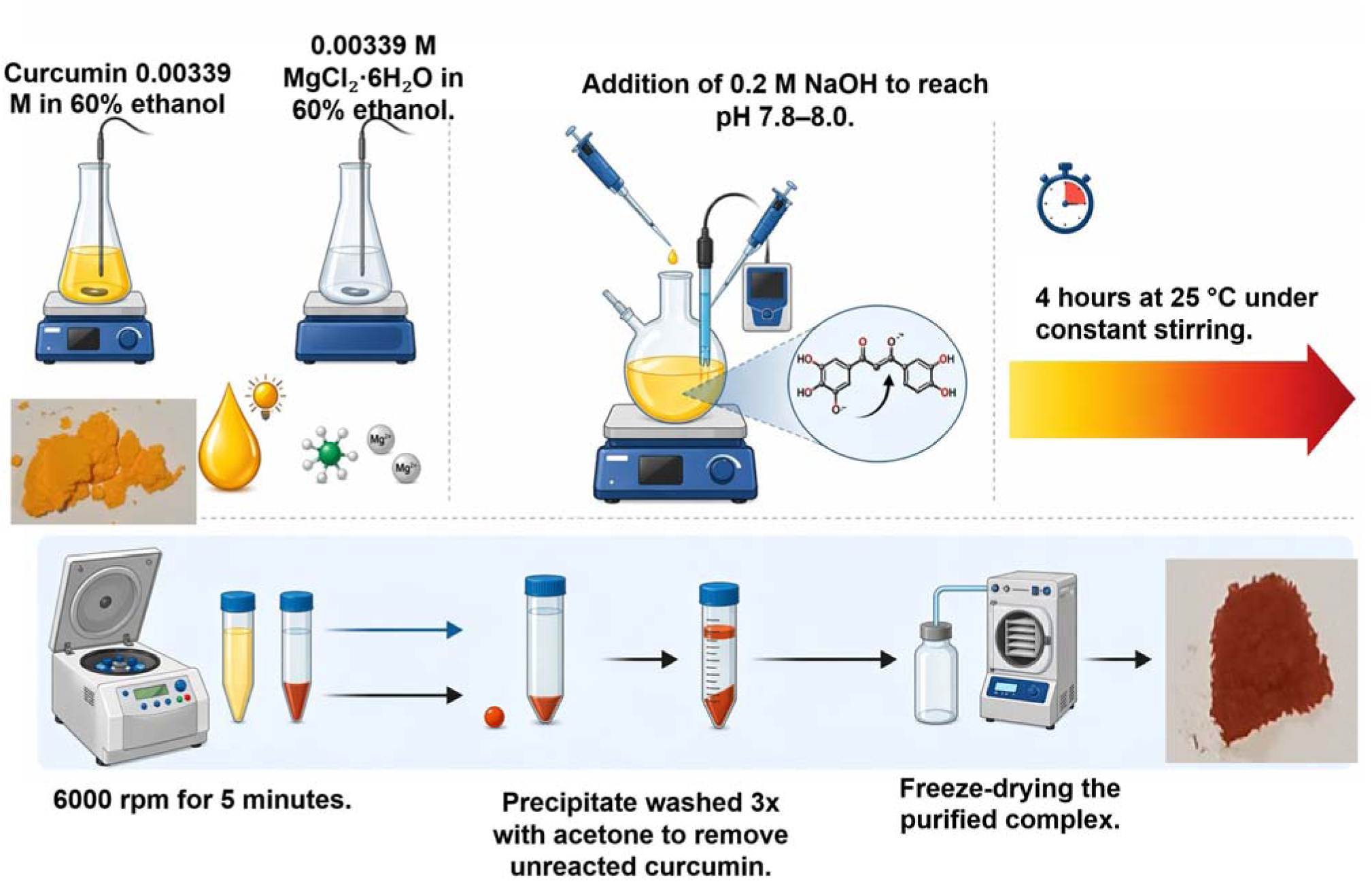
Preparation scheme of Cur-Mg complex

#### 2.1.1 UV–Vis Spectroscopy

UV-Vis spectroscopy indicating changes in the electronic state of curcumin upon interaction with Mg²⁺. Free curcumin exhibited a strong absorption maximum at 430 nm, consistent with the π→π* transition of its conjugated diarylheptanoid chromophore in the enol-keto tautomeric form-a value well within the reported range for curcumin in polar organic solvents (415-435 nm)^45^. The Cur-Mg complex exhibited a pronounced bathochromic (red) shift to 456 nm, accompanied by increased peak intensity, spectral broadening, and an extended absorption tail into the 500–700 nm region.

This 26 nm red shift is characteristic for the metal-ligand coordination Mg²⁺ chelates the central β-diketone oxygens to form a stable six-membered chelate ring, which delocalizes π-electrons and lowers the π→π* transition energy^46^. ligand-to-metal charge-transfer (LMCT) contributions involved in broadening the spectrum. Comparing with literature data confirm that complexation with divalent metals consistently produces red shifts of 5–30 nm with concomitant spectral broadening^47^. These data represent in **Figure 2**. suggest the formation of the Cur-Mg complex and demonstrate stabilization of the curcumin enolate form upon coordination. Curcumin and Cur-Mg comples are viazualized in **Figure 2.(A)** and **Figure 2.(B)**

**Figure. 2:**
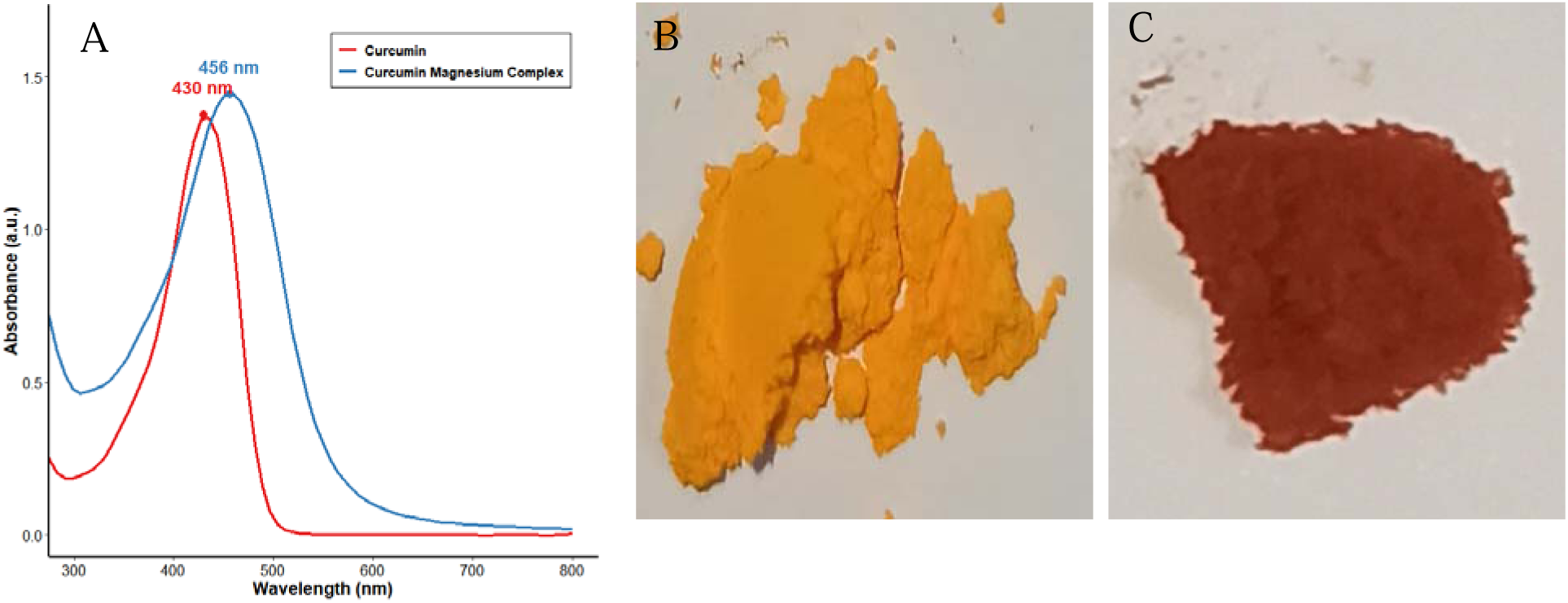
(A) UV–Vis absorption spectra of free curcumin and the Cur-Mg complex. The absorption maximum (λmax) of curcumin appears at 430 nm, while the Cur-Mg complex exhibits a bathochromic shift to 456 nm with spectral broadening, confirming metal-ligand coordination and extended conjugation. (B) Picture of Pure curcumin (C) Picture of Cur-Mg complex

**Figure 3.**
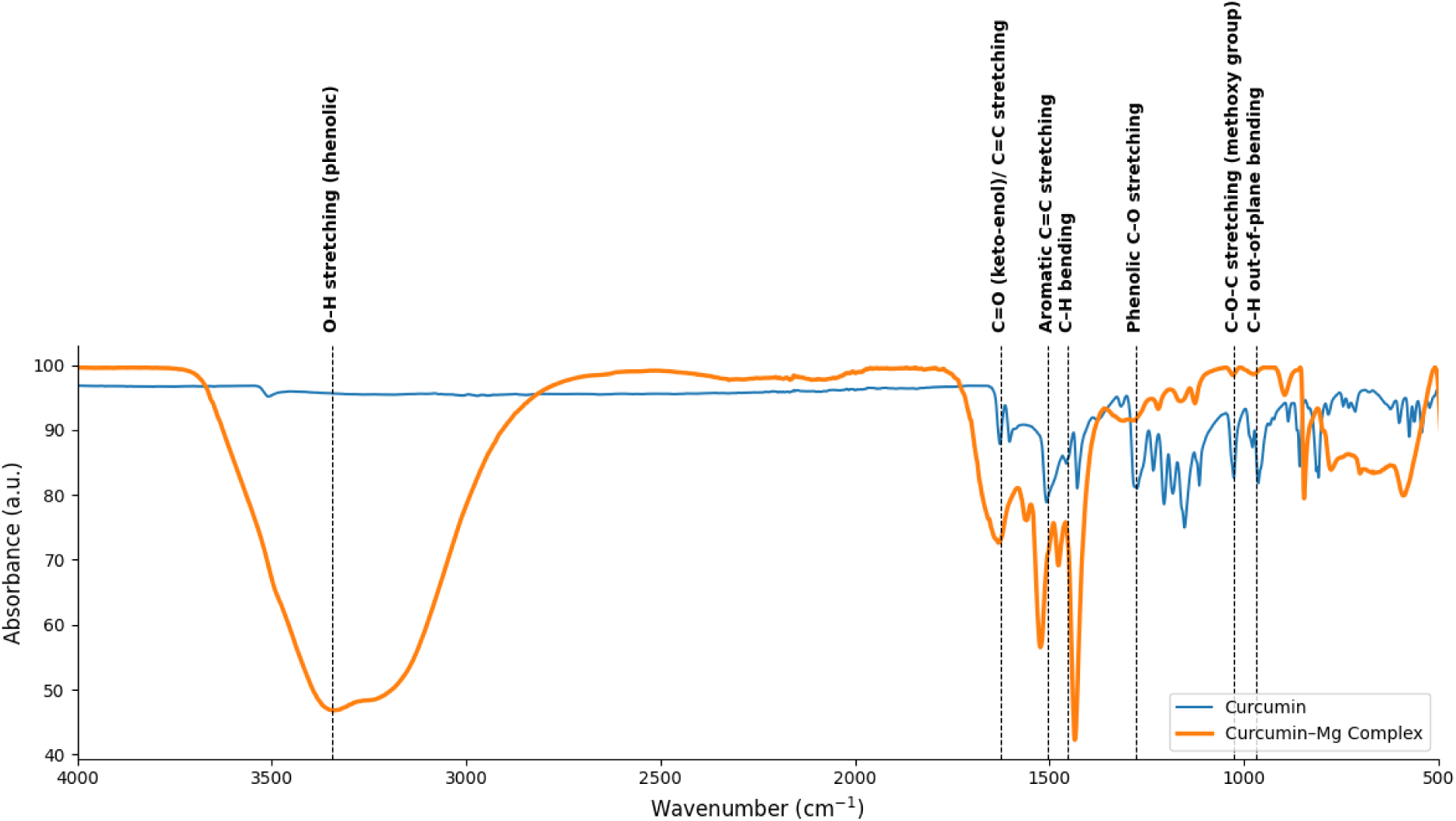
FTIR spectra of curcumin and the curcumin-magnesium (Curcumin-Mg) complex in 500–4000 cm^-1^ range. The spectrum of the Curcumin-Mg complex (orange) clearly shows a broad, sharp band around 3300 cm^-1^ representing the O-H stretching from the phenolic hydroxyl group that is greatly increased compared to the curcumin-Mg complex (blue), indicating participation in the metal complexation. A significant change in the C=O (keto-enol) / C=C stretching band around 1600 cm-1 is seen with increasing intensity and slight frequency shift upon complexation, which proves complexation is likely through the diketone part. Bands due to aromatic C=C stretching and C-H bending (at about 1500-1550 cm-1), phenolic C-O stretching (at about 1270 cm-1) and C-O-C stretching for the methoxy group plus C-H out-of-plane bending (at about 900-1050 cm-1) are conserved in the spectra of curcumin-Mg complex. This confirms that the curcumin framework is maintained. These frequency shifts and differences in the relative band intensities clearly support coordination with Mg through the enolic oxygen and phenolic hydroxyl group of curcumin.

#### 2.1.2 FTIR Spectroscopy

Figure 3. shows an example for comparison, and Table 1 displays the common peaks of the curcumin and Cur-Mg complex. Both FTIR spectra of curcumin and Cur-Mg complexes show characteristic shifts in absorption bands due to coordination of metal-ligand the broad O-H stretching band at ∼3400 cm-1 was significantly broad and its transmittance was decreased greatly (95.7% 49.0%), indicating that OH group is possibly involved in the coordination and enol deprotonation is occurred partially during the chelation. Furthermore, the transmittance of C=O/C=C stretching band of -diketone unit at ∼1628 cm-1 decreased clearly (87.8% 72.1%), which indicates that Mg atom is coordinated via keto-enol system.

**Table 1.** FTIR common peak assignments for curcumin and the Cur-Mg complex.

| Sr.no | Wavenumber | Assignment |
| --- | --- | --- |
| 1 | 3400-3500 | Phenolic O–H stretching |
| 2 | 1628 | C=O / C=C β-diketone |
| 3 | 1510-1515 | Aromatic C=C stretching |
| 4 | 1452 | C–H bending (olefinic) |
| 5 | 1277 | Phenolic C–O stretching |
| 6 | 1150 | C–O–C (methoxy group) |
| 7 | 960-980 | C–H out-of-plane bending |

Changes in the aromatic C=C region (1500-1600 cm) show electron density rearrangement within the conjugated system upon complexation with Mg. Changes in the fingerprint region (1000-1300 cm) that belong to phenolic C-O and methoxy C-O-C vibration suggest interaction between curcumin and Mg. All these spectral shifts can be attributed to chelation of Mg with the -diketone part of the Cur-Mg complex.

#### 2.1.3 X-Ray Diffraction (XRD)

XRD analysis supports complex formation, associated lattice contraction and alternation in crystalline order which can be interpreted by comparing ***Table 2*** and ***Table 3***. As shown in Figure 4A Free curcumin exhibited multiple sharp, high-intensity reflections at 2θ ≈ 14.78°, 17.54°, 18.41°, 21.44°, 23.62°, 24.85°, 25.85°, and 27.64°, consistent with literature^48^. In contrast, the Cur-Mg complex displayed a modified diffraction profile characterised by reduced peak intensities, broadening, and the emergence of new or shifted reflections at lower angles (9.58°, 11.22°, 23.16°, 27.60°, 29.94°, 33.16°, 36.78°, and 40.71°). Visualised in Figure 4B.

**Figure 4.**
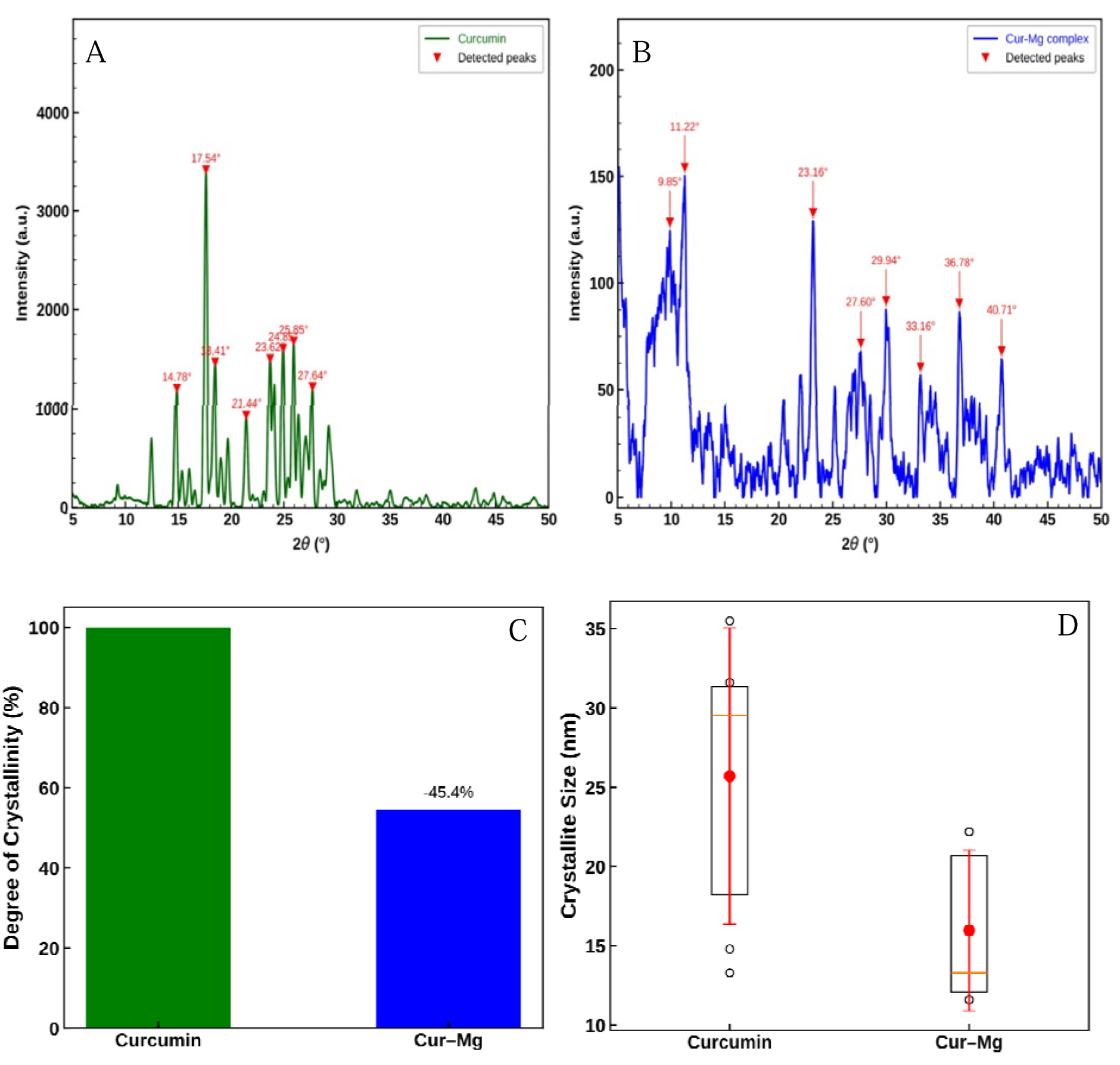
Powder XRD pattern of (A) curcumin, (B) Cur-Mg complex. Red triangles denote the detected diffractions peaks. (C) Quantitative crystallinity (D). Box and whisker of the crystal size. The comparison of the crystallite sizes calculated from the Scherrer equation that decrease from 25.72 ± 9.33 nm for curcumin to 15.98 ± 5.06 nm for Cur-Mg complex; error bars indicate standard deviations. Statistical test has shown a trend of decreasing crystallite sizes, but it was not significantly different (t= 2.198, p=0.0595; one-way ANOVA: F=4.328, p=0.0672).

**Figure 5.**
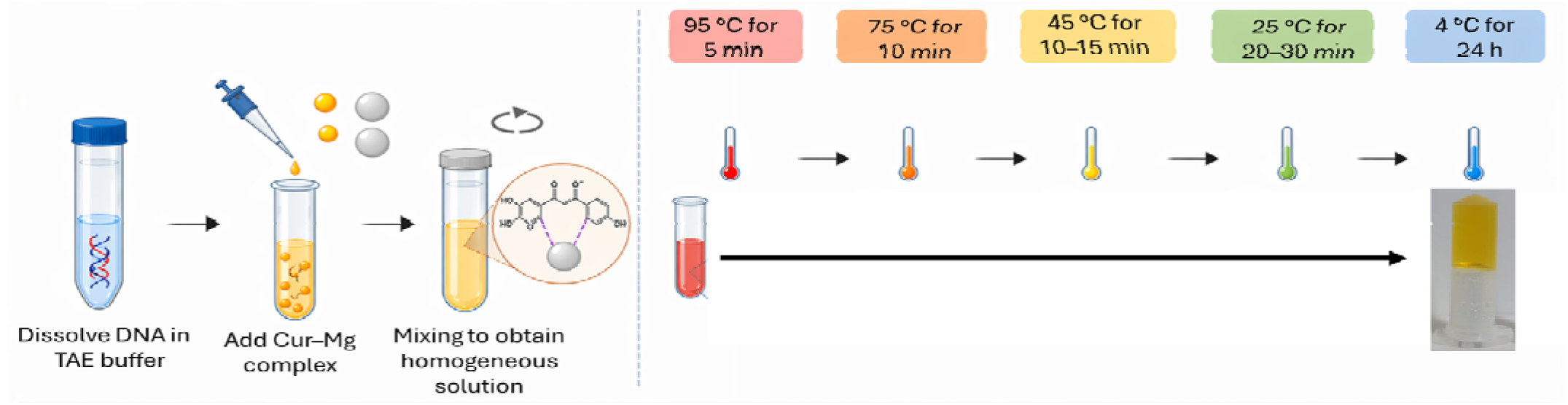
Illustration of the synthesis procedure and thermal gelling of the Cur-Mg-DNA hydrogel. Initially, To make the Cur-Mg-DNA hydrogel genomic DNA was dissolved in the DNA in 1X TAE buffer. Then curcumin-magnesium complex TAE buffer solution was added unto the DNA TAE solution (ratio according to the required formulation). Homogeneous solutions of DNA and Cur_Mg complex was heated at 95 degrees Celsius, for 5 minutes. Then we slowly cooled it down to 75 degrees Celsius for 10 minutes. After that we cooled it down some to 45 degrees Celsius for 10 to 15 minutes. Finally we cooled it down to 25 degrees Celsius for 20 to 30 minutes. This helped the Cur-Mg-DNA hydrogel come together in a controlled way and form a network. Left the Cur-Mg-DNA hydrogel in the cold at 4 degrees Celsius for 24 hours. To make it stable. Final Cur-Mg-DNA hydrogel looks like in **Figure 5**. Where methodology of formulation of Cur-Mg-DNA gel is summarized.

We performed quantitative crystallinity analysis using the trapezoidal integration method which reveals a degree of crystallinity of 54.60%, representing a 45.40% decrease relative to crystalline considering curcumin as a reference^49^ vizualized in Figure 4C. Scherrer analysis suggest a non-significant reduction in average crystallite size from 25.7 ± 9.3 nm to 15.98 ± 5.06 nm curcumin and *Cur*-Mg complex respectively, suggesting nanoscale structuring induced by metal coordination. The presence of prominent low-angle reflections in the complex further suggests increased interplanar spacing and lattice reorganization. This partial amorphization is widely associated with enhanced solubility and bioavailability of curcumin-based systems^50^, and is consistent with previously reported curcumin–metal coordination complexes^38^.

Figure 4D. illustrate the crystallite size range in nm obtained from Scherrer analysis. This plot suggest that the minimum size of crystallite size of curcumin is 14.8nm to 35.5nm while orange dot assigned for mean crystallite size which is around 25-26 nm. Cur-Mg complex exhibits smaller crystallite size ranging from 13.3nm to 20.7nm and mean size is 15.98nm.

Bragg’s law-derived interplanar spacing (d-spacing) values demonstrated a substantial structural variations between the Cur-Mg complex and free curcumin. In keeping with its well-ordered crystalline structure, pure curcumin showed d-spacing values ranging from ∼7.10 Å (at 2θ ≈ 12.46°) to ∼3.05 Å (at 2θ ≈ 29.28°). On the other hand, all the measured diffraction angles, the Cur-Mg complex displayed d-spacing values between ∼3.84 Å and ∼2.22 Å. Localised lattice contraction upon metal coordination is suggested by the complex’s emergence of reflections at higher diffraction angles and correspondingly lower d-spacing values in comparison to many native curcumin peaks. Also, a general trend towards the distribution of d-spacing values in lower range and at higher angles with a structural rearrangement, partial disappearance of long-range ordering, is evident from above results. All these changes collectively establish lattice distortion and reordering due to Mg-coordination, which are also corroborated with a reduction in size and crystallinity. Such a disorder is known to be responsible for partial amorphization and thus to enhance the physicochemical properties like bioavailability, solubility in curcumin formulation. ***Table.2*** and ***Table.3*** contains crystallographic parameters for curcumin and Cur-Mg-complex

**Table 2.** X-ray diffraction (XRD) analysis parameters and crystallite size estimation of the Curcumin. The table summarizes diffraction peak positions (2θ), corresponding Bragg angles (θ in degrees and radians), full width at half maximum (FWHM, β in degrees and radians), cosine of θ, and calculated crystallite size (D, nm) determined using the Scherrer equation. The results indicate crystallite dimensions ranging from 14.8 to 35.5 nm.

| Peak<br>$2\theta$ ° | $\theta$ ° | $\theta$ (rad) | FWHM $\beta$<br>(°) | FWHM<br>$\beta$ (rad) | cos $\theta$ | D-spacing<br>(Å) | D (nm) |
| --- | --- | --- | --- | --- | --- | --- | --- |
| 12.462 | 6.231 | 0.10875 | 0.235 | 0.00411 | 0.99409 | 7.102554 | **35.5** |
| 14.782 | 7.391 | 0.12899 | 0.274 | 0.00478 | 0.99169 | 5.98981 | **30.6** |
| 17.541 | 8.771 | 0.15308 | 0.266 | 0.00464 | 0.98831 | 5.052224 | **31.6** |
| 21.447 | 10.723 | 0.18716 | 0.296 | 0.00517 | 0.98254 | 4.138342 | **28.5** |
| 23.765 | 11.882 | 0.20739 | 0.639 | 0.01116 | 0.97857 | 3.741785 | **13.3** |
| 29.277 | 14.639 | 0.25549 | 0.580 | 0.01013 | 0.96754 | 3.047238 | **14.8** |

**Table 3.** X-ray diffraction (XRD) peak parameters and crystallite size analysis of the Cur-Mg complex. The table presents diffraction angles (2θ), corresponding Bragg angles (θ in degrees and radians), full width at half maximum (FWHM, θ), cosine of θ, and calculated crystallite size (D, nm) using the Scherrer equation. The estimated crystallite sizes range from ∼11.6 to 22.2 nm.

| Peak<br>$2\theta$ (°) | $\theta$ (°) | $\theta$ (rad) | FWHM<br>$\beta$ (°) | $\beta$ (rad) | cos $\theta$ | D-spacing<br>(Å) | D (nm) |
| --- | --- | --- | --- | --- | --- | --- | --- |
| 23.167 | 11.583 | 0.20217 | 0.410 | 0.00716 | 0.97963 | 3.83672<br>5 | **20.7** |
| 25.185 | 12.593 | 0.21978 | 0.384 | 0.00670 | 0.97594 | 3.53239<br>3 | **22.2** |
| 30.019 | 15.010 | 0.26197 | 0.712 | 0.01242 | 0.96588 | 2.97425<br>4 | **12.1** |
| 36.931 | 18.466 | 0.32229 | 0.753 | 0.01313 | 0.94851 | 2.43168<br>8 | **11.6** |
| 40.682 | 20.341 | 0.35502 | 0.666 | 0.01162 | 0.93764 | 2.21652<br>1 | **13.3** |

### 2.2 Synthesis and Characterization of Cur-Mg Complex–Loaded DNA Hydrogels

#### 2.2.1 Topology of Cur-Mg–Hgel (Atomic Force Microscopy)

The topographical data obtained from AFM indicate that there is a distinct and structured variation in surface topology depending on the DNA hydrogel concentration and on the treatment with Cur-Mg. As anticipated, on going along each sequence with increment of the Cur-Mg concentration, there is a general increase in the surface roughness as observed with an increasing RMS and Ra values, suggesting increased roughness of the surface at nanoscale level. The greatest roughness values were observed for the 1.5% DNA hydrogel at the highest Cur-Mg concentration (C200). This may suggest a synergetic mechanism in that denser DNA network can better accommodate or accumulate the Cur-Mg complex and hence induce more irregularity on the surface. A relatively lower roughness at low DNA concentrations (e.g. 0.5% and 1%) reflects that at these cases, the matrix is more porous and the structure perturbation upon incorporation of the drug is relatively small. Further, with increasing concentrations, both the Rt (Total Roughness) and the growing distance between minimum and maximum values clearly demonstrate increased topographic irregularities with extension on the vertical axis. It can thus be inferred that both the DNA concentration and the metal-drug complex loading play essential roles in determining the surface nanostructure. Figure 6. compiles 2D/3D AFM Representative images, height distribution histograms, and line profiles for hydrogels with varying DNA concentrations (0.5%, 1%, 1.5%) and Cur–Mg loading (C0–C200). Increasing DNA content and Cur–Mg incorporation leads to enhanced surface roughness, broader height distributions, and more pronounced peak–valley features, indicating increased nanoscale heterogeneity. AFM images of (B1) unloaded DNA hydrogel and (B2) Cur-Mg complex–loaded DNA hydrogel (Cur-Mg–Hgel). Scale bar: 20 um. Insets show height profiles across representative network fibers ***Table. 4*** serves detail parameters of AFM of hydrogel. These analysis demonstrate that both DNA hydrogel concentration D and cur-Mg complex influences the surface morphology and nanoscale roughness. Among the unloaded hydrogels 1_C0 (1% DNA without Cur-Mg complex) posses the smoothest and most uniform surface with lowest RMS value 203.30nm, while increase in DNA concentration i.e 1.5_C0 (1.5% without any Cur-Mg complex) shows enhanced surface irregularities and topological variation which can be the sign of enhanced network aggregation. Incorporation of Cur-Mg complex in gel results in enhanced roughness ad heterogeneity, particularly in 1.5_C200 (1.5% DNA with 200ug/ml Cur-Mg complex) with highest value of RMS which is 682.16 nm with peak to valley hight 6241.15nm. These findings can be evident that Cur-Mg-complex incorporation promotes localized cross linking and structural changes within hydrogel matric leads to progressively rough and more heterogeneous surface architectures.

**Figure 6.**
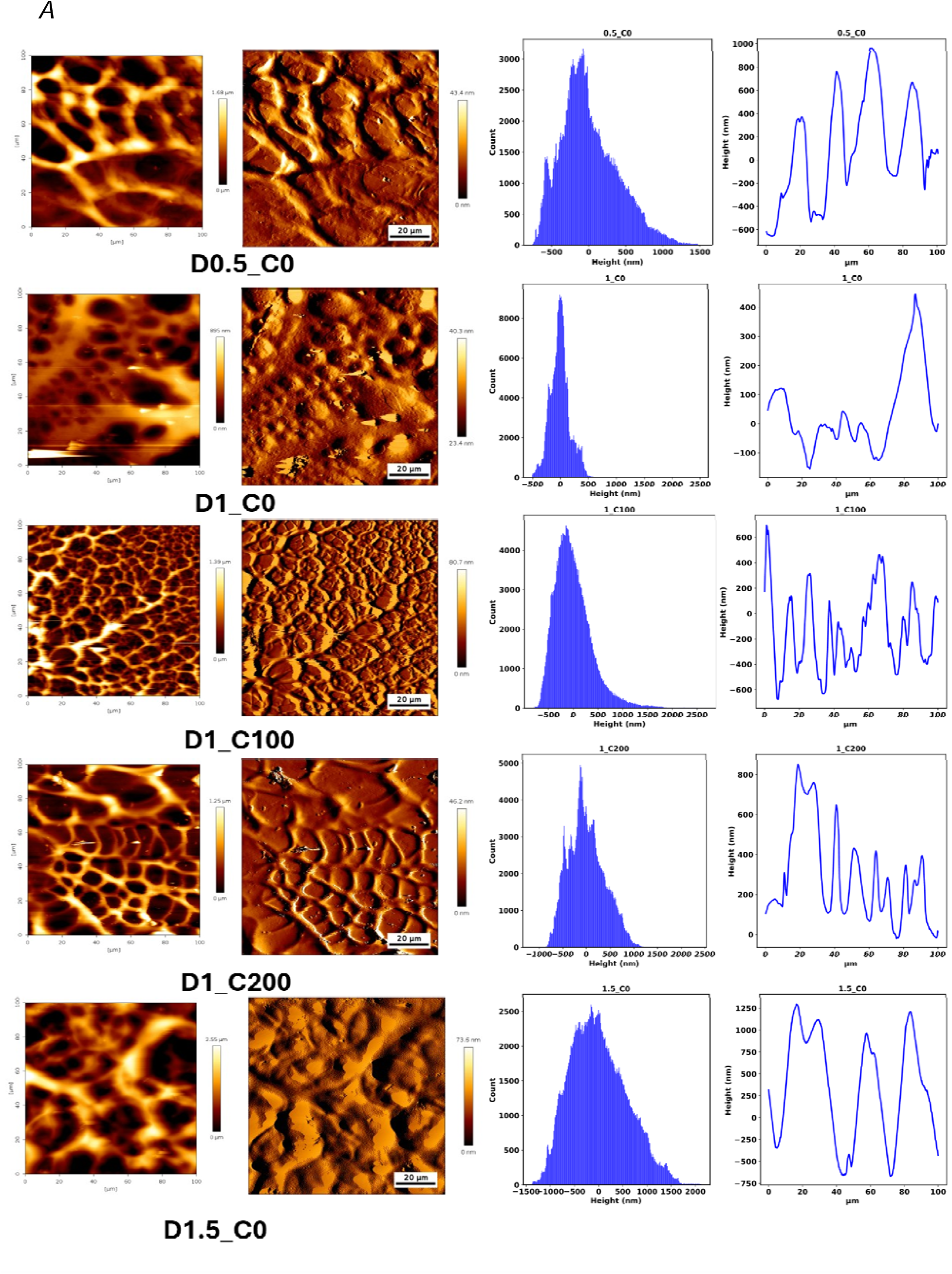
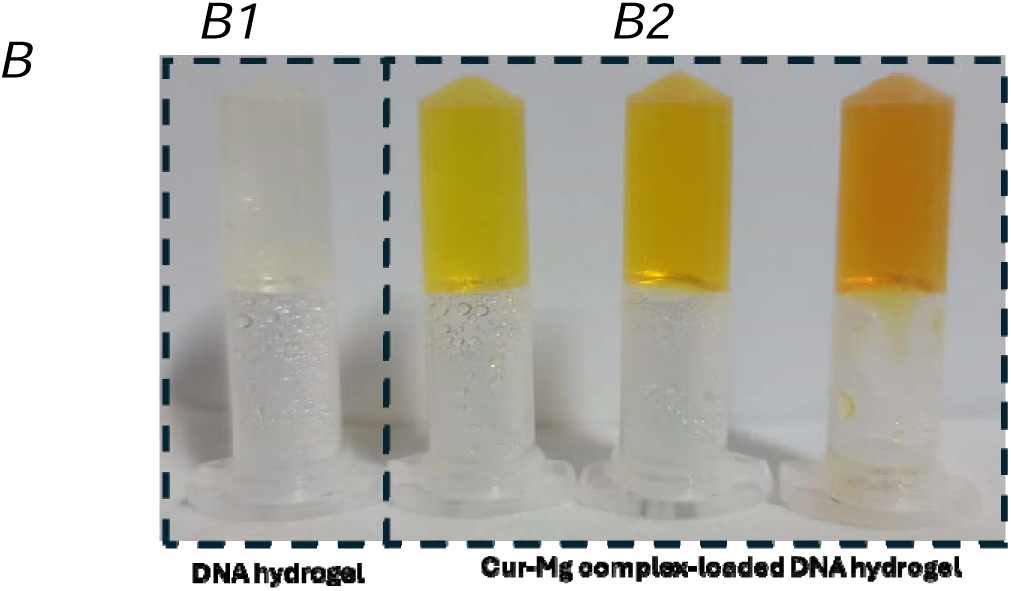
(A) Representative 2D/3D AFM images, height distribution histograms, and line profiles for hydrogels with varying DNA concentrations (0.5%, 1%, 1.5%) and Cur–Mg loading (C0–C200). Increasing DNA content and Cur–Mg incorporation leads to enhanced surface roughness, broader height distributions, and more pronounced peak–valley features, indicating increased nanoscale heterogeneity. AFM images of (B1) unloaded DNA hydrogel and (B2) Cur-Mg complex–loaded DNA hydrogel (Cur-Mg–Hgel). Scale bar: 20 um. Insets show height profiles across representative network fibers.

**Table 4:** Quantitative AFM investigation of surface roughness and topography parameters of DNA hydrogels with Cur-Mg complex.

| **Formulations** | **RMS roughness nm** | **Average roughness nm** | **Peak to valley nm** | **Min height nm** | **Max height nm** |
| --- | --- | --- | --- | --- | --- |
| 0.5_C0 | 382.0667 | 304.3727 | 2312.068 | -765.569 | 1546.5 |
| 1.5_C0 | 579.535 | 470.2651 | 3510.092 | -1382.23 | 2127.862 |
| 1.5_C200 | 682.1637 | 480.5466 | 6241.146 | -2186.9 | 4054.248 |
| 1_C0 | 203.2957 | 141.2222 | 3001.623 | -511.26 | 2490.364 |
| 1_C100 | 384.4197 | 289.4039 | 3516.175 | -812.337 | 2703.838 |
| 1_C200 | 379.3615 | 305.6219 | 3526.668 | -1169.6 | 2357.067 |

#### 2.2.2 FTIR Characterization of Cur-Mg–Hgel

The FTIR spectra of the DNA-Cur-Mg complex hydrogel suggests a complex mixed-mode of interaction with direct coordination to the nucleobases as well as stabilizing interactions with the phosphate backbone. FTIR spectra of DNA and DNA-Cur-Mg gel are presented in Figure 7 while ***Table 5***. contains peak assign for the DNA with their molecular origin. The complex shows the strong change at the guanine-cytosine (G/C) ring vibrational mode, downshifted to 1680 cm and dramatically decrease in intensity from the original position at 1707 cm. This diagnostic band strongly suggests high-affinity binding. High affinity binding interactions to DNA nucleobases can be attributed to direct coordination with the N7 of the purine or with the carbonyl O atoms of the base pairs, as well as to intercalative pi-stacking which stiffens the base pair and enhances binding. In addition, the phosphate backbone undergoes two forms of perturbation. First, there is an upward shift of the asymmetric stretch of the phosphate backbone to 1226 cm (cm), which reflects water molecules being replaced from the DNA hydration shell. Second, the symmetric stretch of the phosphate backbone downshifts to 1070 cm (cm), indicating direct electrostatic interaction of the DNA backbone with the cationic portion of the complex. Despite the strong local perturbations to the DNA scaffold within the complex hydrogel, the DNA appears to maintain its original physiological B-DNA structure with no evidence of transition to the A-form DNA (around 810 cm) by retention of the C2’-endo sugar pucker marker at roughly 831-834 cm. FTIR Peaks of DNA without Cur-Mg complex are approximately similar to the peaks studies in a research article^51^.

**Figure.7:**
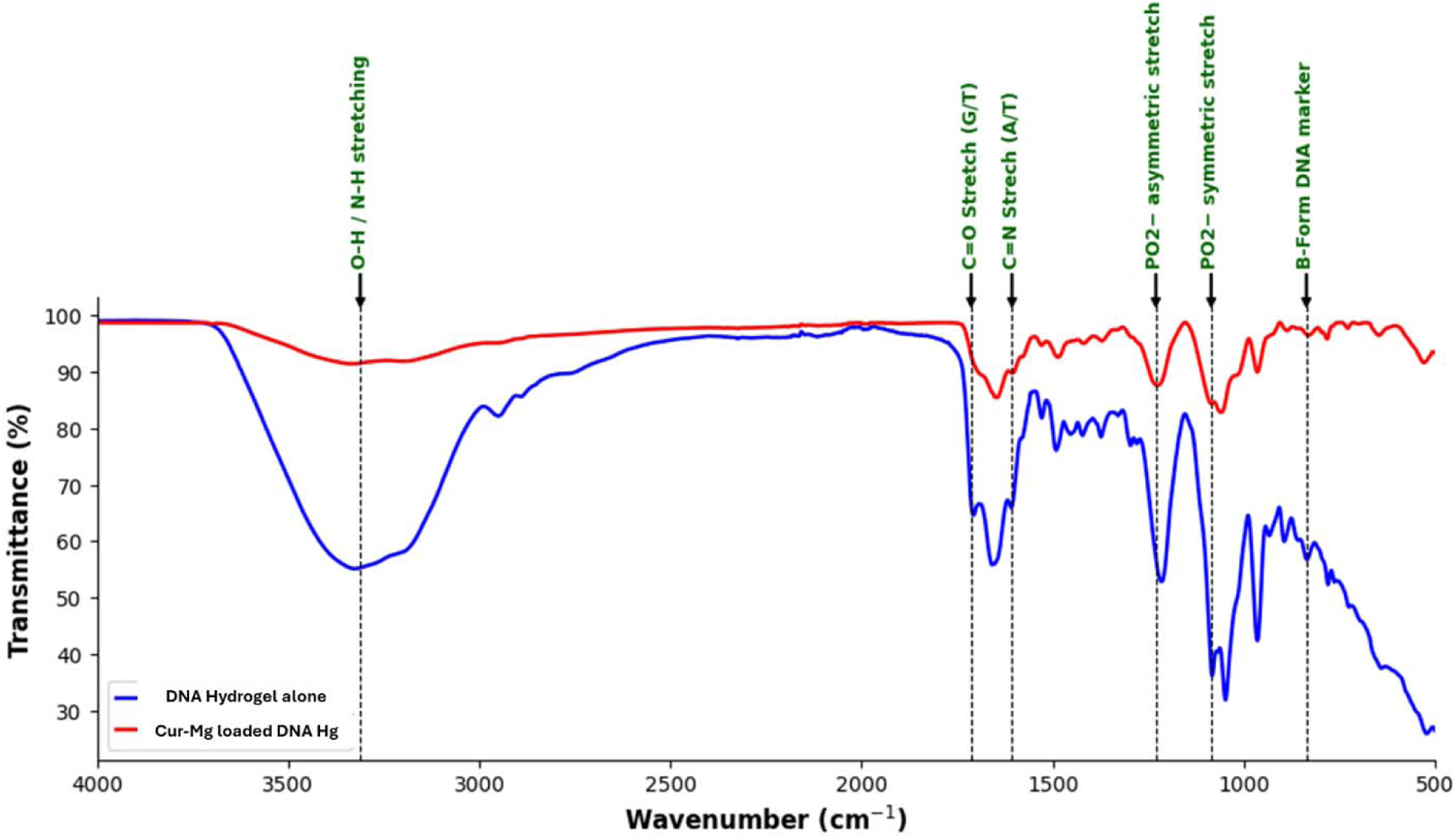
Fourier Transform Infrared (FTIR) spectroscopic analysis of DNA hydrogel alone and curcumin-magnesium complex-loaded DNA hydrogel (Cur-Mg loaded DNA Hg).

**Table 5.** FTIR peak assignments for DNA.

| Wavenumber ($\text{cm}^{-1}$) | Functional Group / Vibration | Molecular Origin |
| --- | --- | --- |
| 3400 | O–H / N–H stretching | Hydroxyl groups (deoxyribose) and nucleobase N–H |
| 1700 | C=O stretching | Carbonyl groups of nucleobases (thymine, guanine) |
| 1600 | C=N stretching | Nucleobase ring vibrations |
| 1230 | $\text{PO}_2^-$ asymmetric stretching | Phosphodiester backbone of DNA |
| 1080 | $\text{PO}_2^-$ symmetric stretching | Sugar–phosphate backbone |
| 800 | B-form DNA marker (C–H / backbone vibration) | Characteristic DNA helical structure |

These spectral changes confirm effective integration of the Cur-Mg complex within the DNA hydrogel matrix without disruption of the fundamental DNA nanostructure.

#### 2.2.3 Mechanical and Rheological Properties

By employing oscillatory shear rheology, we have demonstrated a concentration-dependent increase in the stability of the viscoelastic network of the DNA hydrogel upon addition of curcumin-Mg complex into the DNA hydrogel framework. When examined, neat DNA hydrogel (1.5% w/v, 0 g/mL complex) possesses a G’ (34.4 Pa) and G” (3.71 Pa) (tan 0.108), implying it’s a weakly crosslinked, entanglement-dominated viscoelastic network. By adding 100 g/mL curcumin-Mg complex to the network, G’ increases to 38.8 Pa and tan drops to 0.131, which suggests increased elastic predominance and decreased viscous dissipation as shown in Figure 8. This can be understood through considering the role of curcumin-Mg complex as a supramolecular crosslinking agent; it serves as crosslinker between individual DNA strands by coordinating with metal ion, which causes metal ions or stacking between curcumin ring and DNA base, therefore strengthening the effective crosslinking density of DNA hydrogel network.

**Figure 8:**
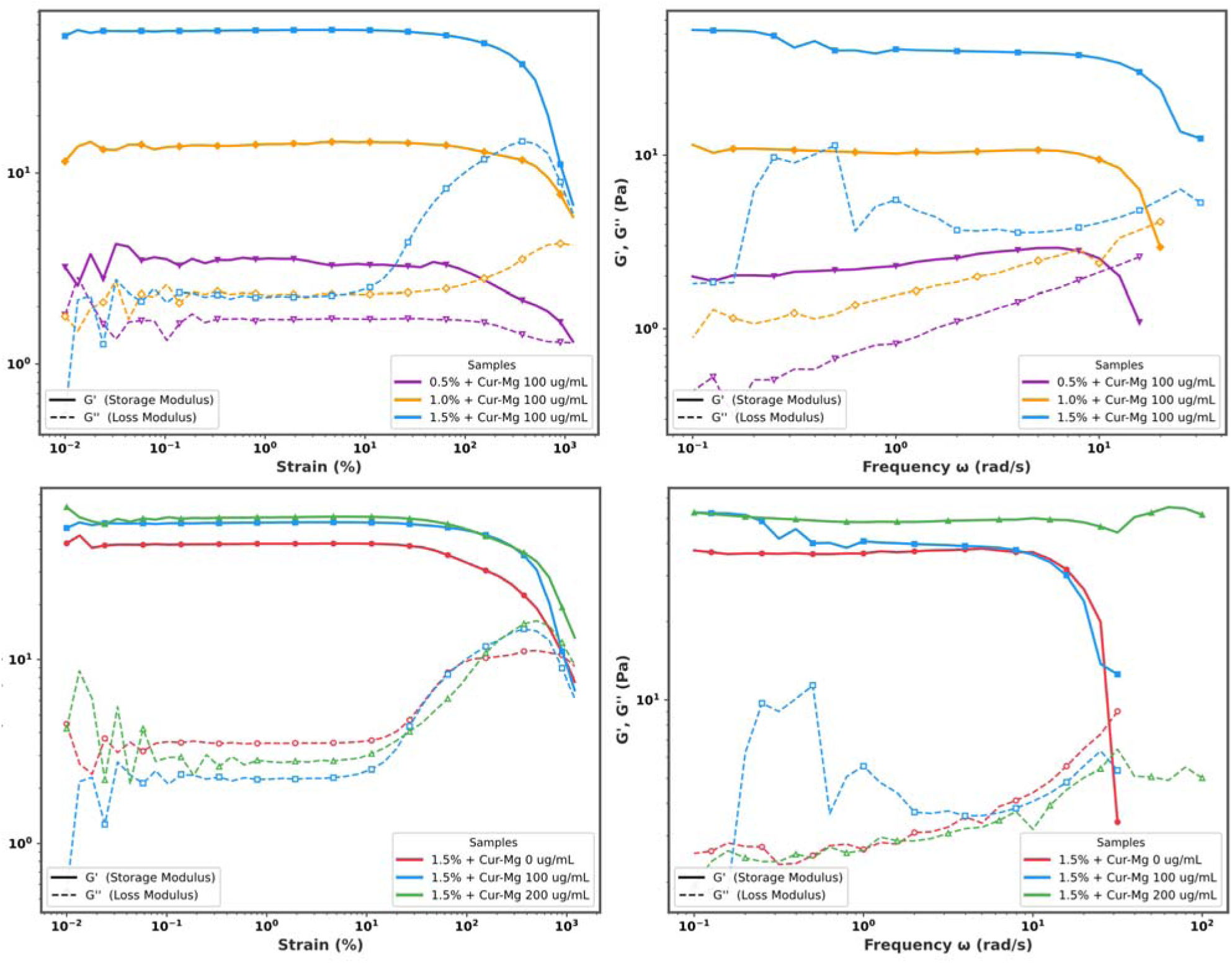
Rheological evaluation of DNA hydrogels: (A) Amplitude sweeps of various concentrations of DNA hydrogels. (B) Amplitude sweep showing difference between Cur-Mg complex loading at 1.5% DNA concentrations. (C) Frequency sweep showing that all formulations are gel like (G > G).

**Figure 9.**
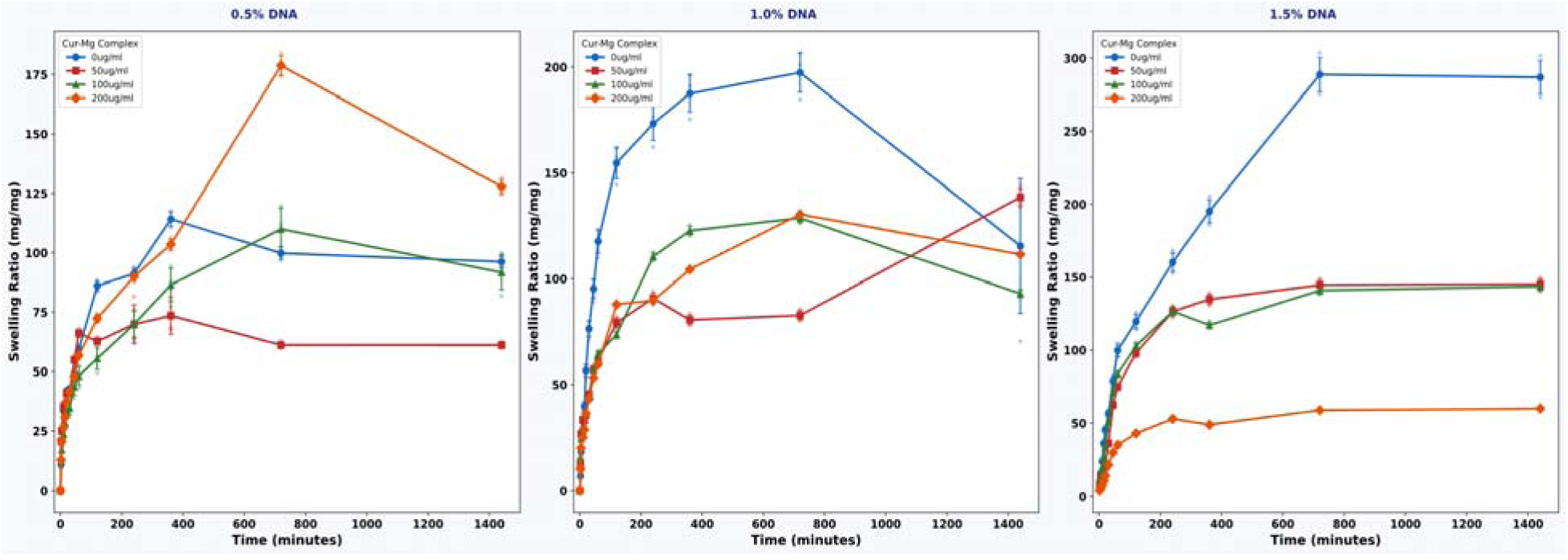
Swelling kinetics of Cur-Mg–Hgel. (A) Time-dependent swelling ratio. Data represent mean ± SD (n = 3).

**Figure 10:**
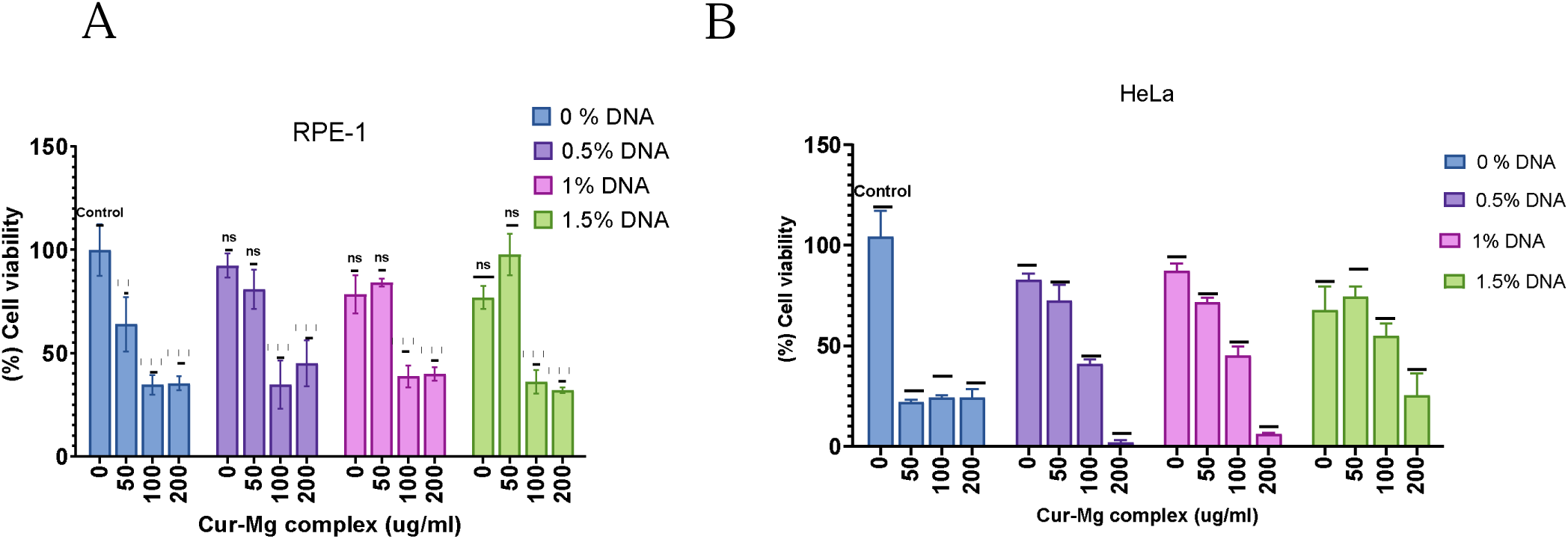
Cell viability of (A) RPE-1 and (B) HeLa cells following treatment with Cur-Mgcomplex at varying DNA concentrations. Percentage cell viability was assessed following treatment with Cur-Mgcomplex (0, 50, 100, and 200 µg/mL) with 0%, 0.5%, 1%, and 1.5% DNA. Two-way ANOVA revealed that Cur-Mgcomplex concentration was the dominant source of variation in both RPE-1 (83.77%, F(3,32) = 135.8, p < 0.0001) and HeLa (73.16%, F(3,32) = 283.2, p < 0.0001) cells. A significant interaction between Cur-Mgcomplex and DNA concentration was observed in both cell lines (RPE-1: F(9,32) = 4.887, p = 0.0004; HeLa: F(9,32) = 28.35, p < 0.0001), while DNA concentration alone was a significant main effect only in HeLa cells (F(3,32) = 8.188, p = 0.0003). Post hoc comparisons (Dunnett’s test) were performed against the untreated control. Data are presented as mean ± SD (n = 3). *p < 0.05, **p < 0.01, ***p < 0.001, ****p < 0.0001; ns = not significant.

At the highest complex loading (200 µg/mL), G′ reached a maximum of 49.8 Pa which is ∼45% increase relative to the control with G″ simultaneously decreasing to 3.53 Pa and tan δ reaching its minimum value of 0.071. The near frequency-independent plateau of G′ across the linear viscoelastic region (LVR) observed for D1.5%_C200 is a hallmark of a percolated, solid-like gel network, wherein the Cur-Mg complex densifies the network architecture and reduces the characteristic relaxation time of the system. The consistently low tan δ values (< 0.1) for D1.5%_C200 confirm the transition to a predominantly elastic hydrogel, in accordance with the theoretical expectation that plateau modulus scales with crosslink density in affine network models.

**Figure 11.**
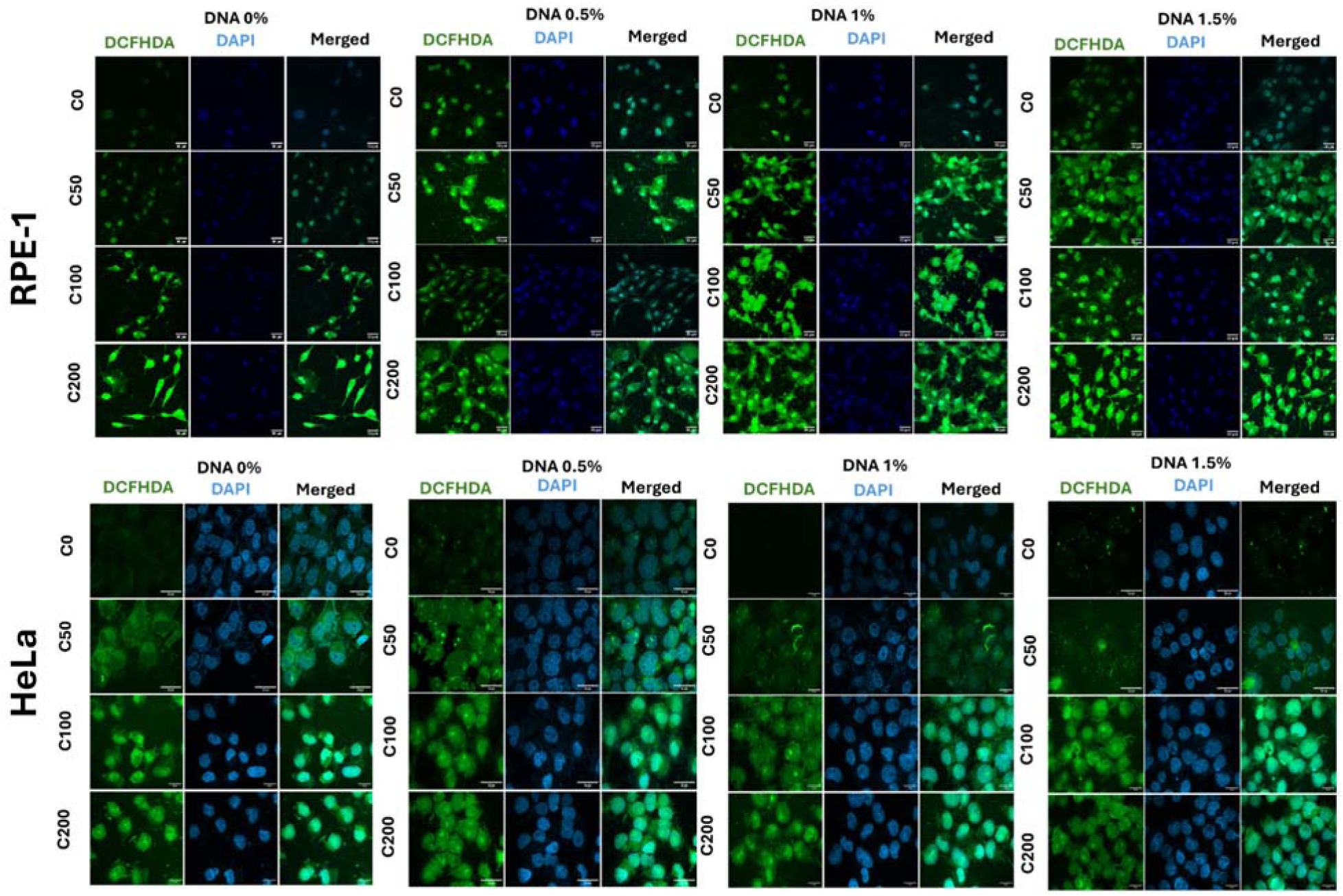
Intracellular reactive oxygen species (ROS) levels in RPE-1 and HeLa cells treated with curcumin-magnesium (Cur-Mg) complex-loaded DNA hydrogels, assessed by DCFH-DA assay. Fluorescence microscopy representative images showing ROS generation in (A) RPE-1 (upper panel) and (B) HeLa (lower panel) cells following 24 h treatment with varying concentrations of Cur-Mg complex (C0 = 0 µg/mL, C50 = 50 µg/mL, C100 = 100 µg/mL, C200 = 200 µg/mL) incorporated into DNA hydrogels at four DNA concentrations (0%, 0.5%, 1%, and 1.5% w/v). Intracellular ROS was detected using the fluorescent probe 2′,7′-dichlorofluorescein diacetate (DCFH-DA; green channel), and nuclei were counterstained with DAPI (blue channel). Merged images represent the overlay of DCFH-DA and DAPI signals. Increased green fluorescence intensity indicates elevated ROS levels. Scale bar = 30 µm

**Figure 12.**
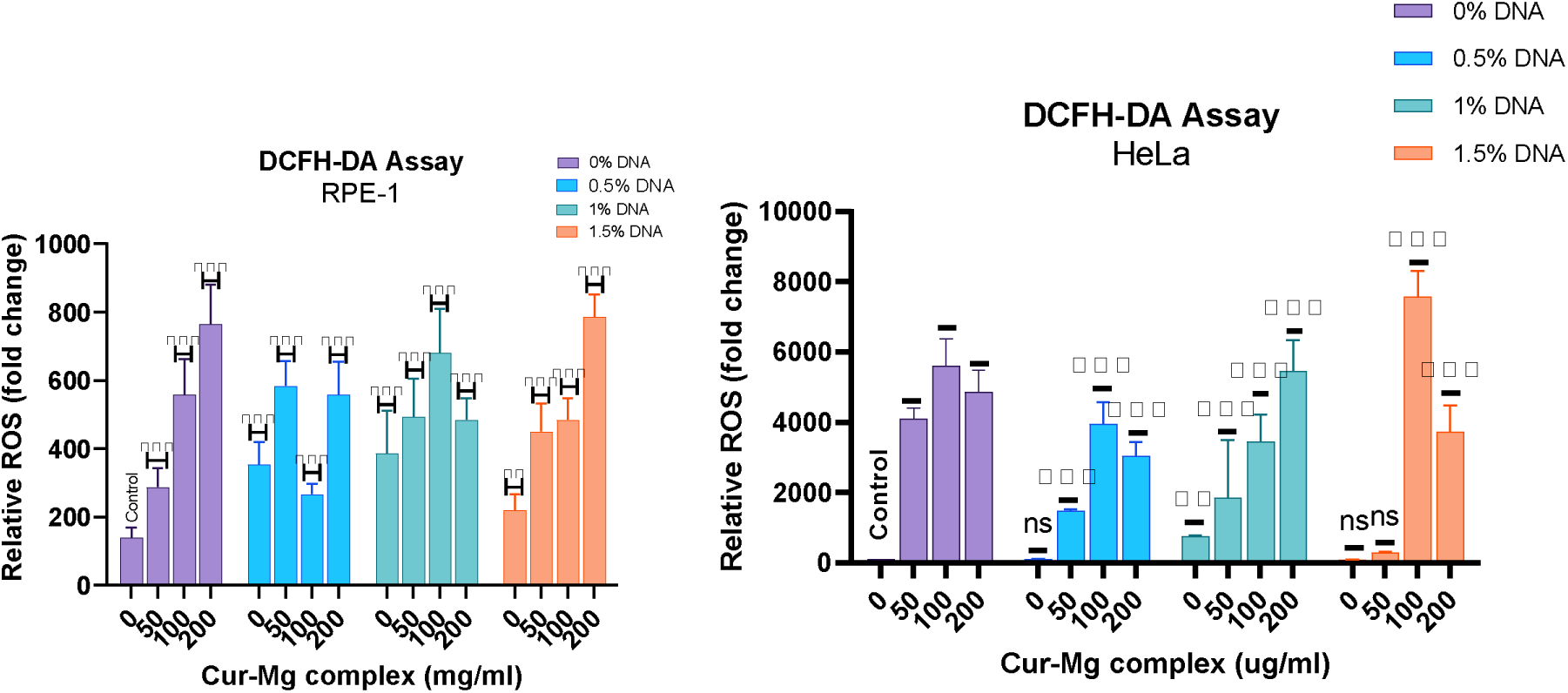
Intracellular ROS generation in HeLa and RPE-1 cells treated with Cur-Mg complex-loaded DNA hydrogels, quantified by DCFH-DA fluorescence assay. Relative intracellular reactive oxygen species (ROS) levels (expressed as fold change in mean fluorescence intensity, MFI) in **(A) RPE-1** (left) and **(B) HeLa** (right) cells following treatment with curcumin-magnesium (Cur-Mg) complex at concentrations of 0, 50, 100, and 200 µg/mL incorporated into DNA hydrogels at 0%, 0.5%, 1%, and 1.5% (w/v) DNA. ROS levels were measured using the 2′,7′-dichlorofluorescein dacetate (DCFH-DA) fluorescence assay. Statistical significance was determined by two-way ANOVA followed by Dunnett’s multiple comparisons test against the untreated control (0 µg/mL Cur-Mg, 0% DNA). Data are presented as mean ± SD (n = 10 cells/ replicate). Significance levels are denoted as **p < 0.01, ****p < 0.0001; ns, not significant.(number of replicates=3)

Conversely, lowering the DNA concentration to 0.5%w/v and keeping the complex concentration constant at 100 ug/mL led to a decrease of G’ to 9.94 Pa and an increase of G (1.94 Pa) and tan (0.195), representing more than two times higher dissipative nature compared to D1.5%C100. This shows that the percolation threshold for the network is exceeded under a critical DNA concentration and that the Cur-Mg complex is not able to overcome low strand densities on its own. The results obtained from the amplitude sweep further support this claim, with the data for D1%C100 showing an earlier and softer yielding regime compared to D1.5%C100 and D1.5%C200 (lower shear stress response for deformation), which reflects the decreased crosslinking density and the more fragile nature of the gel network.

All rheological data together suggests a reinforcing effect of DNA matrix and Cur-Mg complex by working synergistically in the dual component system; DNA matrix provides an essential topological network and skeleton and Cur-Mg complex contributes to increase crosslinking density, prevent strand relaxation and strengthen mechanically. The LVR plateau, a well-defined plateau with the largest G’ (49.8 Pa) and the lowest tan (0.071), can be achieved at1.5%w/v DNA and 200 ug/mL complex.

#### 2.2.4 Swelling Kinetics

The swelling behavior of the hydrogel network influences their kinetics of drug release and therapeutic performance, since the magnitude and speed of water intake is closely correlated to solute diffusion and the degree of network expansion. To investigate the effect of the concentrations of the DNA fiber and Cur-Mg complex, swelling study was carried out in solvent 1X PBS with pH range 7.2-7.4 a time range up to 24 h at 37°C and 5% CO_2_ for all samples.

It was shown that a concentration dependent inverse relation was observed between DNA fiber content and initial water swelling rate at t= 0-10 min for all concentrations of DNA fiber (0.5, 1.0, 1.5 wt%) and Cur-Mg complex concentration (0, 50, 100, 200 ug/mL). The fastest initial water swelling rate was observed at 0.5 wt % DNA (∼5.0%/min) and decreased with the increase of the fiber concentration (3.7%/min and 2.0%/min for 1.0 and 1.5 wt% DNA, respectively). With increasing density of DNA fiber in the network, there is more extensive crosslinking and entangling, which creates more severe steric impediment to solute movement. This leads to less penetrability of the aqueous front into the core of the hydrogel, and thus delay the achievement of the plateau.

To quantify the swelling behaviour in a physically meaningful and analytically tractable framework, experimental data were fitted to a first-order kinetic expression of the form where S(t) is the swelling ratio at time t, S □ is the equilibrium (maximum) swelling ratio, and k is the apparent first-order rate constant^52^. This model is grounded in assumption that the driving force for further water absorption at any instant is proportion the residual swelling capacity of the network which is well-established for describing the swelling of crosslinked biopolymer matrices^53^. The R and RMSE values (0.825-0.991 and 2.71-19.46 mg/mg, respectively) obtained for all twelve formulations were well within acceptable experimental error, thus proving that the first order expression fits reasonably well with the experimental sigmoidal-to-plateau shape of swelling behavior. R values were highest in all of the 1.5 wt% DNA sets, indicating a tighter fiber network adhered most strongly to a monophase plateau model, while the least dense networks (0.5 wt%) had slightly broader residuals which suggest that a second relaxation that was not completely accounted for y a mono-exponential compared to higher concentration sets.

**Figure 13.**
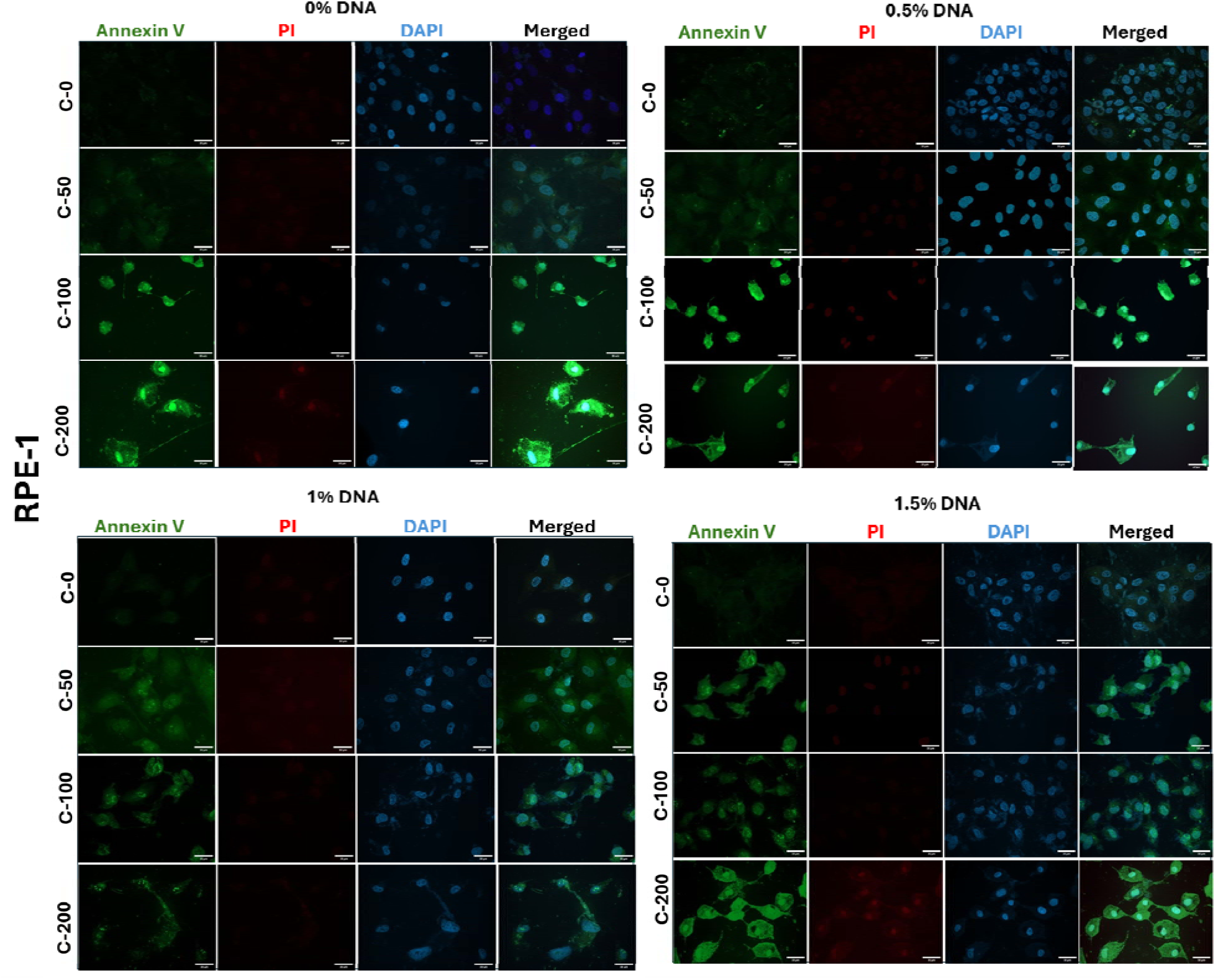
Fluorescence microscopy images of Annexin V/PI staining in RPE-1 cells treated with Cur-Mg complex-loaded DNA hydrogels to assess apoptotic and necrotic cell populations. Representative fluorescence microscopy images depicting apoptosis and cell death in RPE-1 cells following treatment with curcumin–magnesium (Cur-Mg) complex at concentrations of 0 (C-0), 50 (C-50), 100 (C-100), and 200 (C-200) µg/mL incorporated into DNA hydrogels at four DNA concentrations: 0%, 0.5%, 1%, and 1.5% (w/v). Triple-channel staining protocol comprising Annexin V (green), indicative of early and late apoptosis via phosphatidylserine externalisation; propidium iodide (PI; red), marking late apoptotic and necrotic cells with compromised membrane integrity; and DAPI (blue), serving as a nuclear counterstain. Merged images represent the overlay of all three channels. Scale bar = 30 µm.

**Figure 14.**
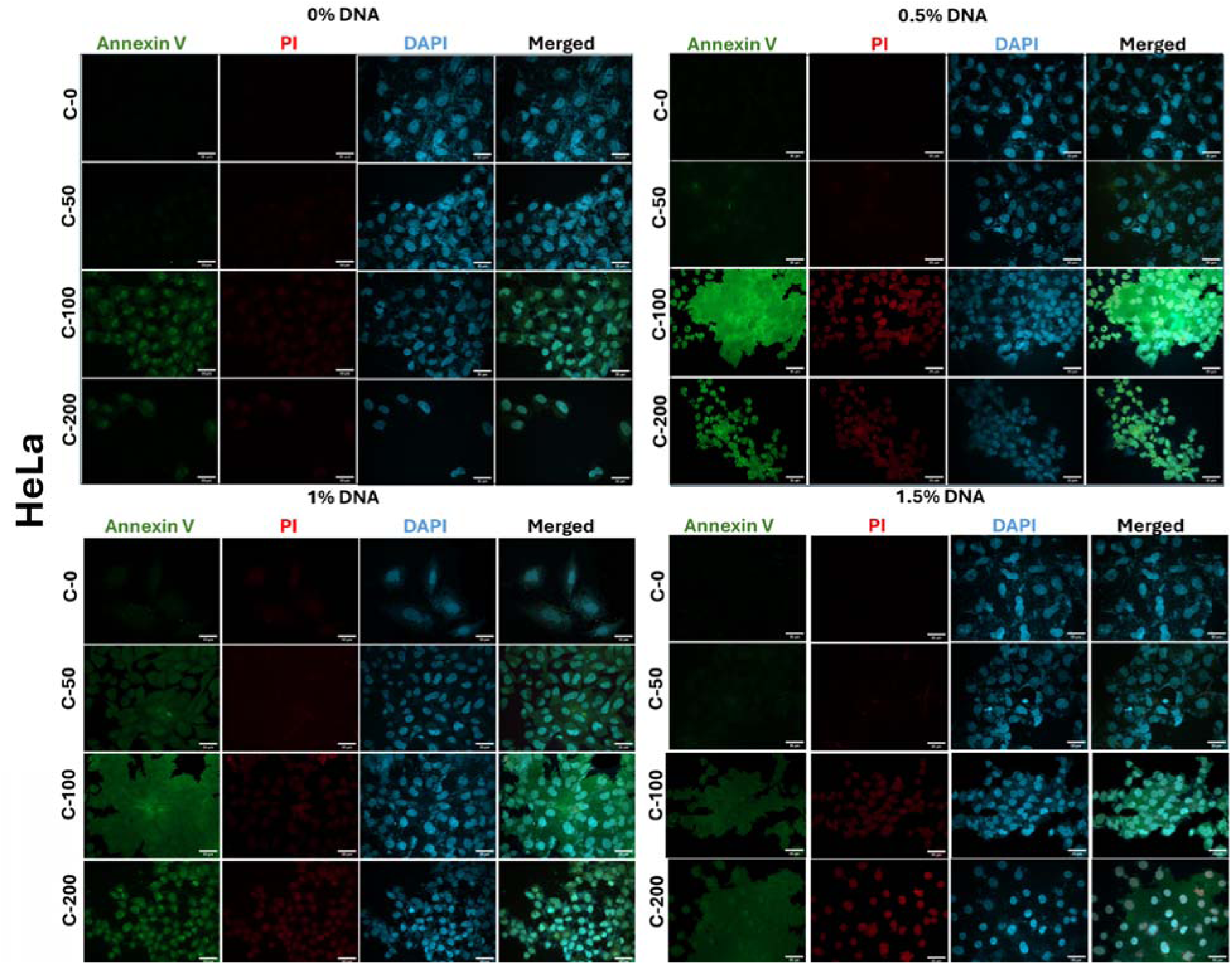
Fluorescence microscopy images of Annexin V/PI staining in HeLa cells treated with Cur-Mg complex-loaded DNA hydrogels to assess apoptotic and necrotic cell populations. Representative fluorescence microscopy images depicting apoptosis and cell death in HeLa cells following treatment with curcumin-magnesium (Cur-Mg) complex at concentrations of 0 (C-0), 50 (C-50), 100 (C-100), and 200 (C-200) µg/mL incorporated into DNA hydrogels at four DNA concentrations: 0%, 0.5%, 1%, and 1.5% (w/v). Triple-channel staining protocol comprising Annexin V (green), indicative of early and late apoptosis via phosphatidylserine externalisation; propidium iodide (PI; red), marking late apoptotic and necrotic cells with compromised membrane integrity; and DAPI (blue), serving as a nuclear counterstain. Merged images represent the overlay of all three channels. Scale bar = 30 µm.

**Figure 15.**
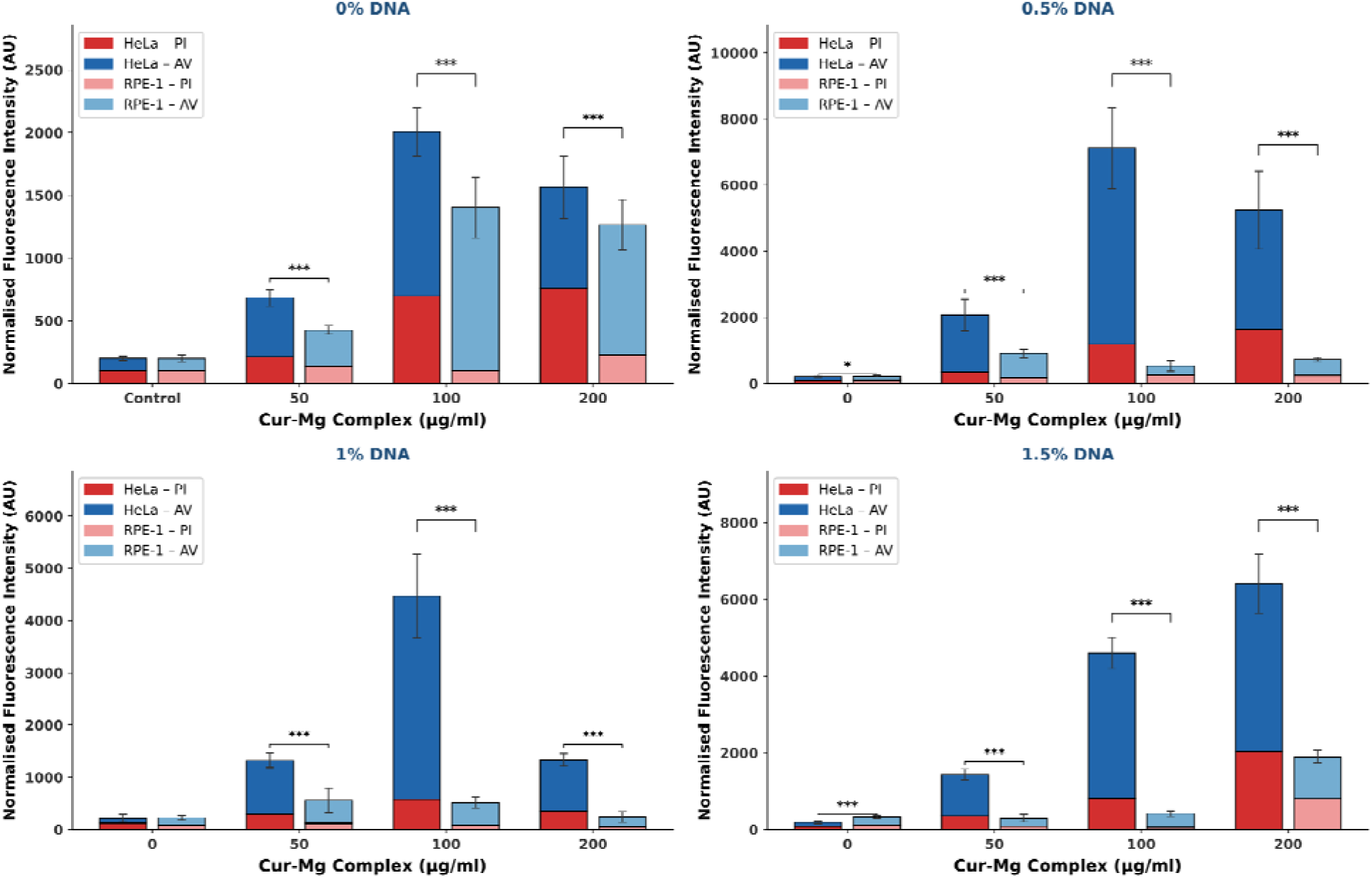
Stacked Annexin V and Propidium Iodide fluorescence in HeLa and RPE-1 cells following Cur-Mg complex treatment. Normalised fluorescence intensities of Annexin V (AV; early apoptosis, upper segment) and Propidium Iodide (PI; late apoptosis, lower segment) are shown for HeLa (dark) and RPE-1 (light) cells treated with 0–200 µg/ml Cur-Mg complex at (A) 0%, (B) 0.5%, (C) 1%, and (D) 1.5% DNA loading. Error bars = SD (n = 10 cells/replicate). stars above individual bars indicate significance vs untreated control by difference between HeLa and RPE-1 at each dose by Tukey’s HSD (three-way ANOVA). * p < 0.05, ** p < 0.01, *** p < 0.001, **** p < 0.0001.(number of replicates=3)

The equilibrium swelling ratio *S* was strongly modulated by both the DNA fibre concentration and Cur-Mg loading. In the absence of Cur-Mg, *S* increased markedly with fibre content: 98.70 ± 5.00 mg/mg at 0.5 wt%, 168.94 ± 9.07 mg/mg at 1.0 wt%, and 281.02 ± 16.45 mg/mg at 1.5 wt%. This positive scaling of equilibrium capacity with fibre density might provide the greater number of hydrophilic binding sites and interchain empty spaces available for water accumulation in more densely fibre-laden networks. The involvement of Cur-Mg produced a concentration-dependent reduction of *S* in most formulations, most favourably at 1.5 wt% DNA where *S* declined from 281.02 ± 16.45 mg/mg (0 µg/mL) to 55.26 ± 1.50 mg/mg at 200 µg/mL Cur-Mg a reduction exceeding 80%. This sudden suppression might be the result of additional ionic and coordination crosslinks formed between magnesium cations and the phosphate backbone of DNA, which tighten the network mesh and restrict chain relaxation supported by FTIR data. similarly the first-order rate possesses *k* exhibited an inverse relationship with *S*: formulations with lower equilibrium capacity tended to display higher *k* values and an e.g., 0.054 min⁻¹ at 0.5 wt% / 50 µg/mL Cur-Mg which reflecting rapid saturation of a smaller swelling reservoir, whereas high-capacity networks (1.5 wt%, 0 µg/mL) swelled slowly (*k* = 0.0045 min⁻¹) as solvent progressively permeated the dense matrix over many hours.

A biphasic swelling trajectory was discernible across all DNA concentrations, characterised by an initial rapid-absorption phase (t = 0–60 min), a transitional plateau region (t = 120–360 min), and a secondary slower-rise phase culminating in a peak at approximately t = 720 min (12 hr) for the majority of formulations. After 12 hr, swelling ratios either stabilised or reduced by small extent by 24 hr, which can be indicative of partial network relaxation, chain disentanglement, or the onset of osmotic re-equilibration between the gel interior and the surrounding medium. This biphasic character was most pronounced in the 1.5% DNA with 0 µg/mL Cur-Mg group, which reached 281.02 ± 16.45 mg/mg at 12 hr before declining marginally at 24 hr, and in the 0.5% DNA groups, which equally peaked near 12 hr. Compared Cur-Mg-crosslinked formulations with 200 µg/mL exhibited a limited swelling capacity and an suppressed secondary phase, suggesting that additional ionic crosslinking not only limits total uptake but also dampens the long-range polymer relaxation responsible for the secondary rise. The time-to-plateau behaviour therefore reflects a competition between osmotic driving force which favours continued uptake and elastic network resistance, with Cur-Mg crosslinking systematically shifting this balance toward earlier and lower saturation.

**Table 6.** Parameters of the first order kinetic swelling studies of DNA-Cur-Mg complex hydrogels under various DNA (0.5%, 1.0%, 1.5% w/v) and Cur-Mg complex (0-200 g/mL) concentrations. Equilibrium swelling capacity (S) and swelling rate constant (k) derived from first order kinetics fitting equation are tabulated along with their regression coefficients of determination (R) and Root Mean Square Error (RMSE) of the fittings. It can be clearly observed that swelling capacity of the hydrogels increases with increasing DNA concentration while the swelling capacity decreases with higher content of Cur-Mgcomplex, while diffusion kinetics also change because the increasing amount of network increases.

| DNA (%) | Cur-Mg ($\mu\text{g/mL}$) | $S_{\infty}$ (mg/mg) | $k$ ($\text{min}^{-1}$) | $R^2$ | RMSE |
| --- | --- | --- | --- | --- | --- |
| 0.5 | 0 | $98.70 \pm 5.00$ | 0.0203 | 0.923 | 9.49 |
| 0.5 | 50 | $64.67 \pm 3.08$ | 0.054 | 0.891 | 6.89 |
| 0.5 | 100 | $88.80 \pm 6.17$ | 0.0143 | 0.877 | 10.9 |
| 0.5 | 200 | $138.50 \pm 12.95$ | 0.007 | 0.853 | 18.29 |
| 1 | 0 | $168.94 \pm 9.07$ | 0.0195 | 0.933 | 17.1 |
| 1 | 50 | $95.65 \pm 7.64$ | 0.0219 | 0.825 | 14.71 |
| 1 | 100 | $111.49 \pm 6.36$ | 0.0155 | 0.917 | 11.45 |
| 1 | 200 | $109.41 \pm 5.14$ | 0.0154 | 0.945 | 9.23 |
| 1.5 | 0 | $281.02 \pm 16.45$ | 0.0045 | 0.957 | 19.46 |
| 1.5 | 50 | $141.45 \pm 2.96$ | 0.0108 | 0.991 | 4.84 |
| 1.5 | 100 | $132.28 \pm 3.60$ | 0.0155 | 0.983 | 6.49 |
| 1.5 | 200 | $55.26 \pm 1.50$ | 0.0155 | 0.983 | 2.71 |

This data is obtained from 60% of the total swelling only because this model assumes a constant, geometry-independent and pure diffusion-controlled mechanism^53^. The power-law model (*S/S = k t*ⁿ) was used to analyze this data. the amount something swells compared to the amount it can swell equals some constant times time raised to a certain power. In this equation the power is called the diffusion exponent. The constant is called the transport rate constant, which is described in the case of the mass-transport mechanism of the mass-transport mechanism.***Table 7***^53^. The value of n helps us understand how transport works: n ≤ 0.5 means diffusion is driven by concentration gradients, 0.5 < n < 1.0 means transport is a mix of diffusion and polymer relaxation n ≥ 1.0 means transport is controlled by relaxation.

**Table 7.** Power-law model (Mt/M∞ = k·tⁿ) parameters described the swelling kinetics of DNA-Cur-Mg complex hydrogels at varying DNA concentrations (0.5%, 1.0%, and 1.5%) and Cur-Mg-complex loadings (0–200 µg/mL). The diffusion exponent (n) which indicates the transport mechanism Fickian to non-Fickian, while the rate constant (k) display swelling kinetics. High R² values demonstrate good agreement between experimental data and model fitting to support the applicability of the power-law model for describing diffusion-controlled swelling behavior in the hydrogel network.

| DNA (%) | Cur-Mg ($\mu\text{g/mL}$) | n (diffusion exponent) | k (rate constant) | $R^2$ |
| --- | --- | --- | --- | --- |
| 0.5 | 0 | 0.442 | 0.0902 | 0.9102 |
| 0.5 | 50 | 0.267 | 0.233 | 0.9283 |
| 0.5 | 100 | 0.382 | 0.0888 | 0.9865 |
| 0.5 | 200 | 0.398 | 0.059 | 0.9943 |
| 1 | 0 | 0.423 | 0.059 | 0.6928 |
| 1 | 50 | 0.306 | 0.1073 | 0.8901 |
| 1 | 100 | 0.396 | 0.09 | 0.9759 |
| 1 | 200 | 0.504 | 0.0597 | 0.9872 |
| 1.5 | 0 | 0.619 | 0.0226 | 0.9816 |
| 1.5 | 50 | 0.642 | 0.0286 | 0.8804 |
| 1.5 | 100 | 0.668 | 0.0345 | 0.9669 |
| 1.5 | 200 | 0.726 | 0.0184 | 0.9721 |

For the 0.5 DNA series n ranged from 0.267 to 0.442. This shows that water transport mostly follows diffusion. At fibre density the network relaxes quickly and molecular diffusion is the slowest step.

The 1.0 wt% series had exponents between 0.306 and 0.504. This indicates a transition to transport especially at 200 µg/mL Cur-Mg. Here n equals 0.504. As crosslink density increases it starts to limit chain mobility. The 1.5 wt% DNA series showed exponents between 0.619 and 0.726. This confirms that in DNA networks with Cur-Mg crosslinks swelling is a result of both solvent diffusion and polymer relaxation working together.

In summary the kinetic parameters and diffusion exponents paint a picture. As DNA fibre content increases, water transport shifts from Fickian, to diffusion. It also increases equilibrium swelling capacity. Slows down the rate constant. Cur-Mg crosslinking works independently to change S and network mesh size. It does not change the transport regime at a given fibre density though.

### 2.3 In Vitro Biological Evaluation

In vitro biological evaluation of Cur-Mg-Hgel was conducted on HeLa a cell of human cervical carcinoma and RPE-1 a cell of Retinal Pigment Epithelial cells - line 1, non-cancerous cell lines. We use three complementary assays (i) the MTT cell viability assay, DCFH-DA assay for intracellular reactive oxygen species (ROS) quantification and Annexin V/Propidium Iodide (PI) dual-staining. Together, these assays provide mechanistic insight into the mode of action of the formulation.

#### 2.3.1 Cell Viability Assessment (MTT Assay)

The viability assay for the curcumin-Mg complex were conducted in both HeLa (cancerous) and RPE-1 (non-cancerous) cell lines using a MTT assay which forms crystal as amount of mitochondrial dehydrogenase. Two-way ANOVA was performed to assess the independent and interactive effects of Cur-Mg complex concentration (0, 50, 100, and 200 µg/mL) and DNA concentration (0%, 0.5%, 1%, and 1.5%) on cell viability.

In HeLa cells, Cur-Mg complex concentration was the dominant factor compare to the RPE-1 cells, as vizualized in ***Figure.10 (B)*** accounting for minimum formazan crystal formation compare to the control, followed by a significant interaction effect between Cur-Mg complex and DNA concentration and a comparatively smaller but significant contribution from DNA concentration alone.

According to Post hoc analysis (Dunnett’s test) all treatment groups exhibited significantly reduced viability compared to the untreated control. A pronounced dose-dependent reduction in HeLa cell viability was observed with increasing Cur-Mg-Complex concentrations, with the most toxic effect at 200 µg/mL, where viability dropped at 0.5%, 1%, and 1.5% DNA concentrations, respectively. While 0% DNA with 200 µg/mL retained viability of approximately 24.40%, while the addition of DNA - particularly at 0.5% and 1% further potentiated cytotoxicity, suggesting a synergistic interaction between the curcumin-Mg complex and DNA in HeLa cells.

RPE-1 cells line show comparatively a distinct response profile. Cur-Mg complex concentration remained the primary driver of toxicity however, DNA concentration alone had no statistically significant main effect. A significant interaction was nonetheless detected,indicating context-dependent modulation by DNA. Post hoc analysis showed that at 0 and 50 µg/mL Cur-Mg complex with all DNA concentrations show no significant differences compared to the untreated control, but Cur–Mg complex alone group at 50 µg/mL show significant cytotoxicity but this toxixity is comparatively less than 50ug/ml alone in HeLa cells. Due to absence of DNA Cur-Mg complex dose of 100 and 200 µg/mL, all groups showed highly significant reductions in viability in both RPE-1 and HeLa however, viability values remained comparatively higher than in HeLa cells at equivalent concentrations, ranging from approximately 32% to 46% at 200 µg/mL. surprisingly, at 50 µg/mL, RPE-1 cells in the 1% and 1.5% DNA groups maintained viability above 84% and 97%, respectively, suggesting a degree of selectivity of the Cur-Mgcomplex toward cancerous cells at lower concentrations.

All together, these findings reavel the Cur-Mg complex exerts potent, concentration-dependent cytotoxicity that is significantly more pronounced in HeLa cancer cells than in non-cancerous RPE-1 cells. The significant interaction between Cur-Mg complex and DNA concentration in HeLa cells, but not in RPE-1 cells, further suggests a selective mechanism of action that may involve DNA-mediated enhancement of cytotoxicity in cancer cells. These results support the potential of curcumin-based complexes as selective anticancer agents. And hydrogel with 0.5% DNA and 50ug/ml cur-Mg complex can be the better dose compared to others

#### 2.3.2 Intracellular Reactive Oxygen Species Quantification (DCFH-DA Assay)

To evaluate the oxidative stress-inducing potential of the Cur-Mg complex loaded DNA hydrogels in cancer and non-cancer cell lines, intracellular reactive oxygen species (ROS) levels were measured using the DCFH-DA fluorescence assay. A two-way ANOVA was performed to get the independent and combined effects of Cur-Mg complex concentration (0, 50, 100, and 200 µg/ml) and DNA concentration (0%, 0.5%, 1%, and 1.5%) on ROS generation in both HeLa and RPE-1 cells.

In HeLa cells, the Cur-Mg complex concentration was identified as the primary source of variation in ROS levels. DNA concentration contributed a smaller but statistically significant proportion of variance and a significant interaction between the two factors was also observed, indicating that the effect of DNA concentration on ROS was dependent on the level of Cur-Mg complex present. Dunnett’s multiple comparisons test revealed that the majority of treatment conditions produced significantly elevated ROS compared to the untreated control (0 µg/ml Cur-Mg, 0% DNA). Notably, treatment with Cur-Mg complex at 100 µg/ml combined with 1.5% DNA resulted in the greatest increase in ROS, followed by 200 µg/ml Cur-Mg with 1% DNA.

In the absence of Cur-Mg complex, DNA alone at 0.5% and 1.5% concentrations did not produce a statistically significant elevation in ROS, suggesting that the oxidative response in HeLa cells is primarily driven by the Cur-Mg complex rather than the DNA carrier alone.

In RPE-1 cells, a broadly similar pattern was observed, though with notable differences in the relative contribution of each factor. The Cur-Mg complex remained the dominant driver of ROS variation, while the interaction term accounted for a proportionally larger share of variance compared to HeLa cells, suggesting greater synergy between Cur-Mg and DNA concentrations in this cell line. DNA concentration alone also reached statistical significance. Strikingly, unlike HeLa cells, all 15 pairwise comparisons against the untreated control were statistically significant in RPE-1 cells, including conditions where DNA was applied without any Cur-Mg complex. This indicates that RPE-1 cells exhibit heightened basal sensitivity to oxidative stress, even in response to the DNA carrier vehicle alone. The largest absolute ROS increase in RPE-1 cells was observed at 200 µg/ml Cur-Mg without additional DNA, with further significant elevations across all higher Cur-Mg doses.

Despite this heightened sensitivity, the overall ROS induction remained substantially more pronounced in HeLa cells under combined treatment conditions, supporting the concept of enhanced oxidative stress in cancer cells^54^. Collectively, these findings indicate that the Cur-Mg-DNA system enables ROS generation through both concentration-dependent and interaction-driven mechanisms, highlighting its potential for inducing oxidative stress-mediated cytotoxicity while emphasizing the importance of optimizing conditions to achieve a favorable therapeutic balance between cancer and normal cells.

#### 2.3.3 Apoptosis Analysis

The apoptotic effects of the Cur-Mg complex were assessed using Annexin V (early apoptosis) and Propidium Iodide (PI; late apoptosis) fluorescence in HeLa and RPE-1 cells. Two-way ANOVA revealed that both Cur-Mg concentration and DNA loading percentage significantly influenced apoptosis in both cell lines, with a significant interaction between the two factors indicating that DNA formulation percentage modulates the apoptotic response to the drug. In HeLa cells, Annexin V and PI fluorescence increased progressively with Cur-Mg dose, with Cur-Mg concentration alone accounting for over 50% of total variance in both stains, confirming it as the primary driver of cell death. Post-hoc analysis showed significant apoptotic induction from 50 µg/ml onward, peaking at 200 µg/ml with 0.5% DNA for Annexin V and at 200 µg/ml with 1.5% DNA for PI. RPE-1 cells, while showing statistically significant overall effects, displayed a notably weaker and less consistent dose-dependent response, with several dose DNA combinations failing to reach significance in post-hoc comparisons. Three-way ANOVA directly comparing both cell lines confirmed that HeLa cells underwent significantly greater apoptosis than RPE-1 cells for both Annexin V (Δ = 1429.4 AU, p < 0.001, η²p = 0.884) and PI fluorescence (Δ = 472.5 AU, p < 0.001, η²p = 0.830). The significant CellLine × Curcumin interactions for both markers (η²p = 0.858 and 0.780, respectively) indicate that the Cur-Mg complex preferentially induces apoptosis in cancerous HeLa cells over non-cancerous RPE-1 cells, suggesting selective anticancer activity. The higher Annexin V relative to PI signal across most conditions further suggests that early apoptosis is the predominant mechanism of Cur-Mg-induced cell death.

To further characterise the mode of cell death, the ratio of Annexin V to Propidium Iodide (AV:PI) fluorescence was calculated for each treatment condition using intensity values pre-normalised to the untreated control visualized in Figure 16. In HeLa cells, the AV/PI ratio increased substantially with Cur-Mg treatment peaking at 100 µg/ml across multiple DNA loading conditions (highest at 1% DNA, AV:PI = 6.66), indicating that early apoptosis was the most possible mode of cell death at intermediate concentrations. At 200 µg/ml, the ratio declined compared to the 100 µg/ml peak across most DNA conditions, suggesting a partial shift toward late apoptosis or secondary necrosis at higher doses. In RPE-1 cells, the AV/PI ratio was markedly more variable and lacked a consistent dose-dependent pattern, with notably high ratios at isolated conditions like at 100 µg/ml with 0% DNA results AV/PI 12.98 driven primarily by minimal PI uptake rather than elevated Annexin V signal, reflecting the overall resistance of non-cancerous cells to membrane permeabilisation. Collectively, these observations suggest that the Cur-Mg complex selectively induces early apoptosis in HeLa cells in a concentration-dependent manner, while RPE-1 cells do not exhibit a coherent apoptotic progression under equivalent conditions compare to HeLa.

**Figure 16.**
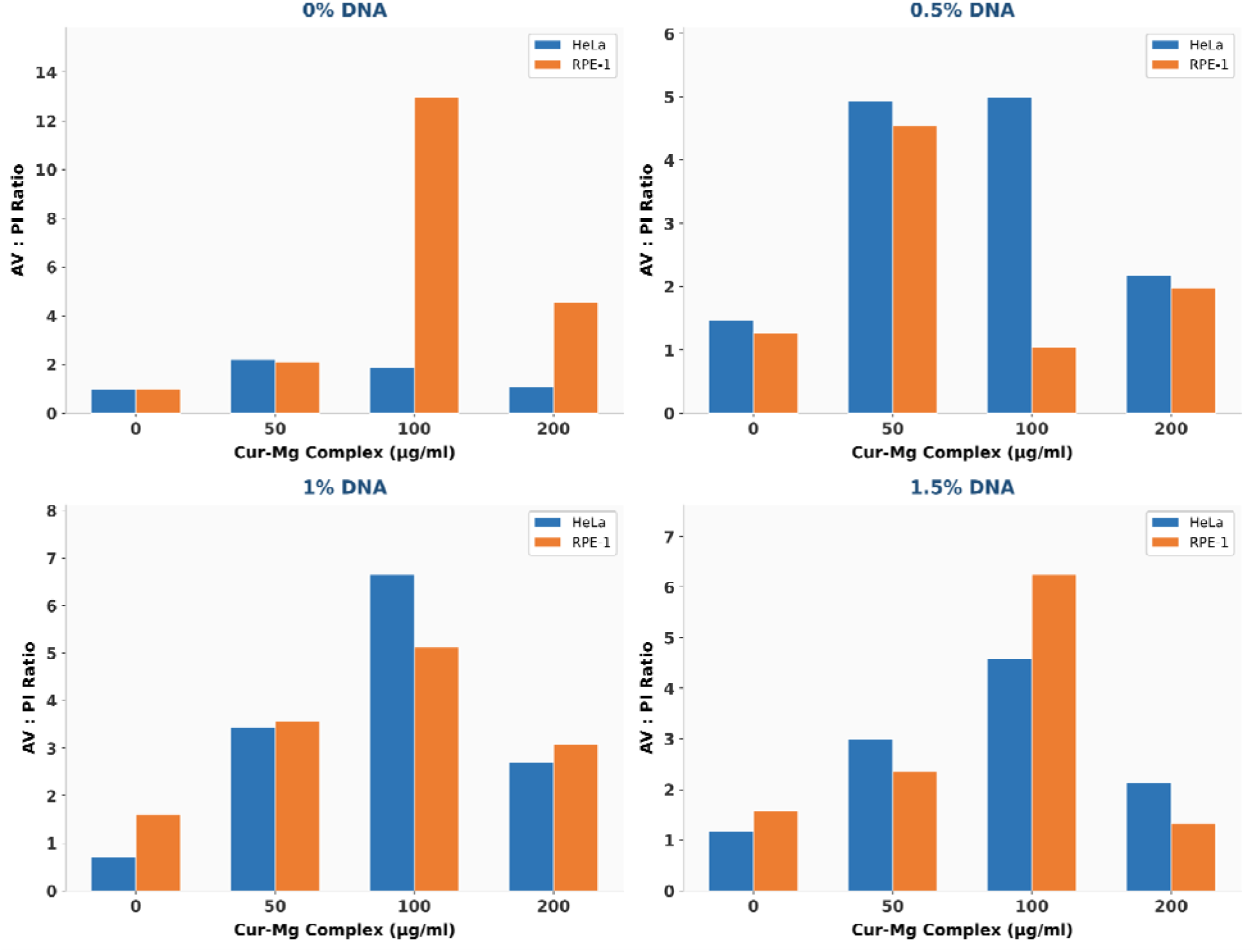
Annexin V to Propidium Iodide (AV:PI) fluorescence ratio in HeLa and RPE-1 cells treated with Cur-Mg complex. The AV:PI ratio was calculated from normalized fluorescence intensities across four Cur-Mg complex concentrations (0, 50, 100, and 200 µg/ml) and four DNA loading percentages (0%, 0.5%, 1%, and 1.5%). Each panel represents a distinct DNA loading condition. Blue bars represent HeLa (cancerous) cells and orange bars represent RPE-1 (non-cancerous retinal epithelial) cellsl. Values above 1 indicate predominance of early apoptosis over late apoptosis/necrosis..

### 2.4 Mechanistic Summary and Synergistic Rationale

Taken together biological assays describe a coupled sequential pathway of Cur-Mg hydrogel induced cancer cell death. ROS accumulation beyond the cytotoxic threshold of HeLa cells which triggers predominantly early apoptosis rather than necrosis. This cascade is selectively amplified in cancer cells because HeLa cells, operating at constituently elevated redox setpoint due to oncogene driven reprograming HeLa posses diminished antioxidant reserve capacity compared to RPE-1, rendering them disproportionately vulnerable to additional pro-oxidant insults.

Concentration dependant network stiffening of hydrogels modulates formulation manner dependent cell death from acute cytotoxicity towards programmed apoptosis at therapeutically relevant dose. Selectivity is most evident at formulation 50ug/ml Cur-Mg complex where 1% DNA hydrogel reduces HeLa viability to 71.64 ±2.23% while preserving RPE-1 viability at 84.08±1.83% a differential absent when the complex is delivered without the matrix and where AV:PI conforms early apoptosis as dominant death mechanism in HeLa but not in RPE-1. These findings validate the formulation D1_C50 is suitable candidate for cancer selectivity.

## 3. Conclusions

A curcumin-magnesium (Cur-Mg) coordination complex was successfully synthesised and thoroughly characterised in this work. It was then integrated into programmable DNA hydrogels (Cur-Mg–Hgel) to provide a unique nanomedicine platform for cancer treatment. Effective metal-ligand coordination, decreased crystallinity, and nanoscale structuring of the complex were confirmed by physicochemical characterisation using UV-Vis spectroscopy, FTIR, and XRD; rheological and swelling analyses showed adjustable mechanical properties and maintained the hydrogel matrix’s structural integrity. Cur-Mg–Hgel’s cancer-selective therapeutic profile was highlighted by an in vitro biological evaluation that demonstrated strong and selective cytotoxicity against HeLa cervical cancer cells with relatively little toxicity towards non-cancerous RPE-1 cells.The system’s controlled and targeted mode of action was highlighted by mechanistic investigations using the DCFH-DA assay, which showed a 3.2-fold increase in intracellular reactive oxygen species (ROS) in treated HeLa cells. Annexin V/PI staining also confirmed that cell death occurred primarily through apoptosis with minimal necrosis. All of these results point to Cur-Mg–Hgel as a biocompatible, pro-apoptotic, and cancer-selective nanomedicine system with significant translational promise. Future research will concentrate on tumor-targeted ligand functionalization to further improve cancer selectivity and cellular uptake, long-term biocompatibility evaluations to guarantee systemic safety, and in vivo pharmacokinetic evaluation to evaluate drug behaviour under physiological situations. These initiatives will be essential to moving the Cur-Mg–Hgel platform from bench-scale research to practical clinical use in cancer treatments.

## 4. Materials and Methods

### 4.1 Materials

Curcumin (≥98% purity, HPLC grade), magnesium chloride hexahydrate (MgCl₂·6H₂O), dimethyl sulfoxide (DMSO), 2′,7′-dichlorofluorescin diacetate (DCFH-DA), and 1,1-diphenyl-2-picrylhydrazyl (DPPH),Salmon Testis Genomic DNA. The MTT reagent kit and Annexin V–FITC/PI apoptosis detection kit were obtained from Thermo Fisher Scientific. HeLa and RPE-1 cells. DMEM, fetal bovine serum (FBS), penicillin/streptomycin solution, and trypsin-EDTA were sourced from HiMedia Laboratories (Mumbai, India). All other chemicals and solvents were of analytical grade and used as received. Milli-Q water (18.2 MΩ·cm) was used throughout.

### 4.2 Synthesis of the Cur-Mg Complex

Curcumin (0.00339 M) was dissolved in 60% ethanol under mild heating (60 °C, 2 min) and protected from light. A 0.00339 M MgCl₂·6H₂O aqueous solution was stirred at 400 rpm, 25 ± 1 °C. The curcumin solution (2 mL) was added dropwise (∼0.2 mL min⁻¹) to the Mg²⁺ solution (10 mL) to achieve a 1:1 molar ratio. The pH was adjusted to 7.8–8.0 using 0.2 M NaOH under continuous stirring. The reaction was maintained for 4 h followed by overnight dark incubation (∼12 h). The complex was isolated by centrifugation (6000 rpm, 5 min), washed three times with acetone, and lyophilized.

### 4.3 Fabrication of Cur-Mg–Loaded DNA Hydrogels

DNA hydrogels were prepared by mixing Genomic salmon testis DNA (at specified DNA concentrations: 0.5%, 1.0%, 1.5% w/v) in 1× TAE/Mg²⁺ buffer (40 mM Tris, 20 mM acetic acid, 1 mM EDTA, 12.5 mM MgCl₂, pH 8.0). The mixtures were heated sequentially from 95 °C for 5 min, 75^0^C for 10min,45^0^C for 10min-15min, 25^0^C for 20-30min and kept in 4^0^C for 24hrs Cur-Mg complex was dissolved in TAE buffer and added at specified concentrations (50,100 and 200 µg/mL) along with the addition of 1× TAE/Mg²⁺ buffer.

### 4.4 Physicochemical Characterization

UV–Vis spectroscopy was performed using a “Thermo Scientific™ BioMate™ 160 UV-Vis Spectrophotometer” over 300–800 nm at 1 nm intervals. FTIR spectra were recorded on a Bruker Tensor 27 ATR-FTIR spectrometer (4000–400 cm⁻¹, 4 cm⁻¹ resolution, 32 scans). XRD patterns were collected using a D8 DISCOVER (Bruker) diffractometer (Cu Kα radiation, λ = 1.5406 Å) over 2θ = 5–90° at 0.02°/step. Crystallite size was calculated by the Scherrer equation and the RCI was determined by trapezoidal integration. AFM imaging was performed on mica substrates using a Bruker Nano wizard Sense AFM. Rheological measurements were conducted on a measured using a stress and strain controlled Modular Compact Rheometer (Anton Paar MCR 302) with a parallel plate geometry at 25 °C. Swelling kinetics were assessed gravimetrically in PBS (pH 7.4, 37 °C). Drug release was quantified by UV–Vis absorbance (430 nm for curcumin) in PBS (pH 7.4)

### 4.5. Swelling Ratio Measurement and Kinetic Analysis

Swelling ratio was measured gravimetrically at predetermined time intervals (2, 4, 10, 15, 20, 30, 45, 60, 120, 240, 360, 720, and 1440 min). Hydrogel samples were removed from the swelling medium (1x PBS, pH 7.4) at each time point, gently blotted with filter paper to remove surface water, and immediately weighed. Swelling ratio (SR) was calculated using the equation: SR = (W_s − W_d) / W_d, where W_s is the swollen weight and W_d is the initial dry weight of the hydrogel, and expressed in mg/mg. Each measurement was performed in triplicate and results are reported as mean ± standard deviation. Time points yielding negative SR values, indicating experimental artefacts such as incomplete surface blotting or sample handling error, were excluded from analysis. Swelling ratio versus time curves were plotted using Google colab with the Matplotlib and NumPy libraries. Mean SR values with ±1 SD shaded bands were plotted for each sample across the full time course (0–1440 min).

To investigate the mechanism of water transport, swelling data were fitted to the Korsmeyer–Peppas power-law model: M_t/M_∞ = k · tⁿ, where M_t/M_∞ is the fractional swelling at time t, k is the swelling rate constant, and n is the diffusion exponent. Data were linearised by plotting log₁₀(M_t/M_∞) against log₁₀(t), and linear regression was performed to determine n and k for each sample and Cur-Mg concentration group. The coefficient of determination (R²) was used to assess goodness of fit. Transport mechanism was classified as Fickian diffusion (n ≤ 0.5), anomalous transport (0.5 < n < 1.0), or Case II transport (n = 1.0) according to established criteria for swelling-controlled systems.

### 4.6 Cell Culture

HeLa and RPE1 cells were sepratly maintained in Dulbecco’s Modified Eagle Medium (DMEM) containing with 10% fetal bovine serum (FBS) and 1% penicillinstreptomycin. Both cells were incubated in T25 flasks at 37°C in incubator which provide a humidified environment containing 5% CO2. The culture medium was replaced as needed, and cells were passaged upon reaching 80–90% confluency. For passaging, spent medium was removed, and cells were treated with 0.25% trypsin EDTA for 5 minutes after 1X phosphate-buffered saline (PBS) which helps to eliminate residual serum, debris, and trypsin inhibitors that could affect enzymatic action. Activity of Trypsin was neutralized by adding 1 mL of complete medium, and the resulting cell suspension was used for either subculturing into new flasks or seeding into well plates, depending on the experimental needs.

### 4.7 MTT Cell Viability Assay

Cell viability was evaluated using the MTT assay in both malignant HeLa and non-malignant RPE-1 cell lines. Cells were cultured in T25 flasks and, upon reaching approximately 80–85% confluency, were trypsinized and seeded into 96-well plates at a density of 2 × 10 cells per well. The cells were allowed to adhere for 24 h at 37 °C in a humidified incubator with 5% CO₂. Following adhesion, the culture medium was replaced with hydrogel leachates prepared by incubating the hydrogels in complete cell culture medium for 24 h. Cells were then exposed to these conditioned media for an additional 24 h under standard culture conditions. After treatment, the medium was carefully removed, and the cells were washed with 1× phosphate-buffered saline (PBS). Subsequently, MTT reagent (0.5 mg/mL; 3-(4,5-dimethylthiazol-2-yl)-2,5-diphenyl tetrazolium bromide) was added to each well and incubated for 4 h at 37 °C to allow the formation of formazan crystals. The MTT solution was then discarded, and 100 μL of dimethyl sulfoxide (DMSO) was added to each well to solubilize the crystals, followed by incubation in the dark for 10 min.Absorbance was measured at 562 nm using a Byonoy Absorbance Microplate Reader. Cell viability was calculated relative to untreated control cells, and the data were analyzed using GraphPad Prism.

### 4.8 Intracellular ROS Quantification (DCFH-DA Assay)

Cells were cultured in T25 flasks and, upon reaching approximately 80–85% confluency, were trypsinized and seeded on coverslips in 24 well plates at a density of 2 × 10 cells coverslip. The cells were allowed to adhere for 24 h at 37 °C in a humidified incubator with 5% CO₂. Following adhesion, the culture medium was replaced with hydrogel leachates prepared by incubating the hydrogels in complete cell culture medium for 24 h. Cells were then exposed to these conditioned media for an additional 24 h under standard culture conditions. After treatment, the medium was carefully removed, and the cells were washed with 1× phosphate-buffered saline (PBS). Treated with 10um DCFHDA solution mixed in serum free media and incubated for 30 min at 37^0^ C, after incubation serum free media was removed and coverslips are washed with 1X PBS 3 times, for fixation 300ul of 4% Para-formaldehyde was added to each well and incubated for 10-15min at room temperature. After 10-15 min 4% Para-Formaldehyde was removed and coverslip was washed with 1x PBS 2 times and mouted on slide with Mowiol containing 1ug/ml DAPI Results are expressed as mean ± SD (n = 10 cells/ replicate).(number of replicates=3)

### 4.9 Apoptosis Analysis (Annexin V–FITC/PI)

Cells were cultured in T25 flasks and, upon reaching approximately 80–85% confluency, were trypsinized and seeded on coverslips in 24 well plates at a density of 2 × 10 cells coverslip. The cells were allowed to adhere for 24 h at 37 °C in a humidified incubator with 5% CO₂. Following adhesion, the culture medium was replaced with hydrogel leachates prepared by incubating the hydrogels in complete cell culture medium for 24 h. Cells were then exposed to these conditioned media for an additional 24 h under standard culture conditions. After treatment, the medium was carefully removed, and the cells were washed with 1× phosphate-buffered saline (PBS) followed by 1X Annexin V binding buffer. Cells were then incubated with Annexin V-FITC and Propidium Iodide (2.5 μL each) in 400 μL of binding buffer for 20 minutes in the dark at room temperature. After incubation, cells were gently washed with 1X PBS and fixed with 2% paraformaldehyde for 15 minutes at 37°C. The fixed cells were washed three times with PBS and mounted onto glass slides using DAPI-containing Mowiol, then left to dry overnight at 4°C. Imaging was performed using a Leica SP8 confocal microscope with a 63X oil immersion objective and appropriate laser settings. The images were processed and analysed using ImageJ software. Results are expressed as mean ± SD (n = 10 cells/ replicate) number of cells.(number of replicates=3)

### 4.10 Relative Crystallinity Index (RCI) Calculation

The RCI was calculated from XRD intensity data (2θ = 5–90°) exported to Microsoft Excel. Baseline correction was applied, and the total area under each diffraction curve was determined by the trapezoidal integration rule: A = Σ[(I + I ₊₁)/2 × (2θ ₊₁ − 2θ)]. Pure curcumin was considered 100% crystalline (reference). RCI (%) = (Area_CurMg / Area_Curcumin) × 100.

### 4.11 Statistical Analysis

All data are presented as mean ± standard error of the mean (SD) from at three independent experiments performed in triplicate. Statistical analyses were performed using GraphPad Prism 9.0. Group comparisons were analyzed by one-way or two-way ANOVA with appropriate post hoc tests (Tukey’s or Bonferroni’s). A p-value of < 0.05 was considered statistically significant.

## Declarations

## Acknowledgements

The authors express their gratitude to every member of the D.B. study group for their helpful criticism and thorough evaluation of the work. We sincerely thank the Indian Institute of Technology Gandhinagar for its financial and infrastructure support. The Central Instrumentation Facility (CIF) at IIT Gandhinagar is acknowledged by the authors for granting access to the characterisation tools utilised in this investigation, including AFM, FTIR, XRD. A.S. thanks financial support from the Prime Minister’s Research Fellowship and the Government of India’s Ministry of Education. The authors thank H.A. for carrying out the rheological tests and supplying the related data, and P.T. for granting access to the rheometer. The confocal microscopy data was obtained by G.P. The Indian National Young Academy of Sciences acknowledges D.B. as a member. J.P and S.B. thanks IIT Gandhinagar’s Department of Biological Sciences and Engineering for providing the resources and research environment needed to complete this work. This work was funded by ANRF-CRG, GSBTM, MoES-STARS and CCRH

## Author Contributions

D.B.: Funding acquisition, writing, review and editing, supervision, resources, and conceptualisation. J.P.: Conceptualisation, research, formal analysis, methodology, first draft writing, and visualisation. S.B.: Research, Approach. A.S.: Data curation, methodology, and investigation. G.P.: Research (collection of image data, confocal microscopy). H.A.: Research, curation of data (rheological experiments). P.T.: Resources (access to rheometers).

## Data Availability

All data supporting the findings of this study are available from the corresponding author upon reasonable request.

